# Amino acid sequence assignment from single molecule peptide sequencing data using a two-stage classifier

**DOI:** 10.1101/2022.09.23.509260

**Authors:** Matthew Beauregard Smith, Zack Booth Simpson, Edward M. Marcotte

## Abstract

We present a machine learning-based interpretive framework (*whatprot*) for analyzing single molecule protein sequencing data produced by fluorosequencing, a recently developed proteomics technology that determines sparse amino acid sequences for many individual peptide molecules in a highly parallelized fashion [1] [2]. Whatprot uses Hidden Markov Models (HMMs) to represent the states of each peptide undergoing the various chemical processes during fluorosequencing, and applies these in a Bayesian classifier, in combination with pre-filtering by a k-Nearest Neighbors (kNN) classifier trained on large volumes of simulated fluorosequencing data. We have found that by combining the HMM based Bayesian classifier with the kNN pre-filter, we are able to retain the benefits of both, achieving both tractable runtimes and acceptable precision and recall for identifying peptides and their parent proteins from complex mixtures, outperforming the capabilities of either classifier on its own. Whatprot’s hybrid kNN-HMM approach enables the efficient interpretation of fluorosequencing data using a full proteome reference database and should now also enable improved sequencing error rate estimates.

## Introduction

Proteins are key components of living organisms, but their heterogenous chemical natures often complicate their biochemical analyses, and consequently, the state of protein identification and quantification methods (e. g., mass spectrometry, antibodies, affinity assays) has generally tended to lag the remarkable progress exhibited by DNA and RNA sequencing technologies. However, improvements to protein analyses could potentially directly inform better biological understanding and better translate into biomedicine and clinical studies. Thus, the field of single molecule protein sequencing attempts to apply concepts from DNA and RNA sequencing to protein analyses in order to take advantage of the high parallelism, sensitivity, and throughput potentially offered by these approaches [3] [4] [5] [6].

Fluorosequencing is one such single-molecule protein sequencing technique inspired by methods used for DNA and RNA [1] [2]. In fluorosequencing, proteins in a biological sample are denatured and cleaved enzymatically into peptides. The researcher then chemically labels specific amino acid types, or alternatively, specific post-translational modifications (PTMs), within each peptide with different fluorescent dyes, then covalently attaches the peptides by their C-termini to the surface of a single-molecule microscope imaging flow-cell (**Figure 1A**). Sequencing proceeds by alternating between acquiring fluorescence microscopy images of the immobilized peptides and performing chemical removal of the N-terminal-most amino acid from each peptide, using the classic Edman degradation chemistry [7] [8] (**Figure 1B**). In this manner, the sequencing cycle (corresponding to amino acid position) at which different fluorescent dyes are removed is measured on a molecule-by-molecule basis, with these data collected in parallel for all the peptide molecules observed in the experiment (**Figure 1C**).

**Figure 1.**
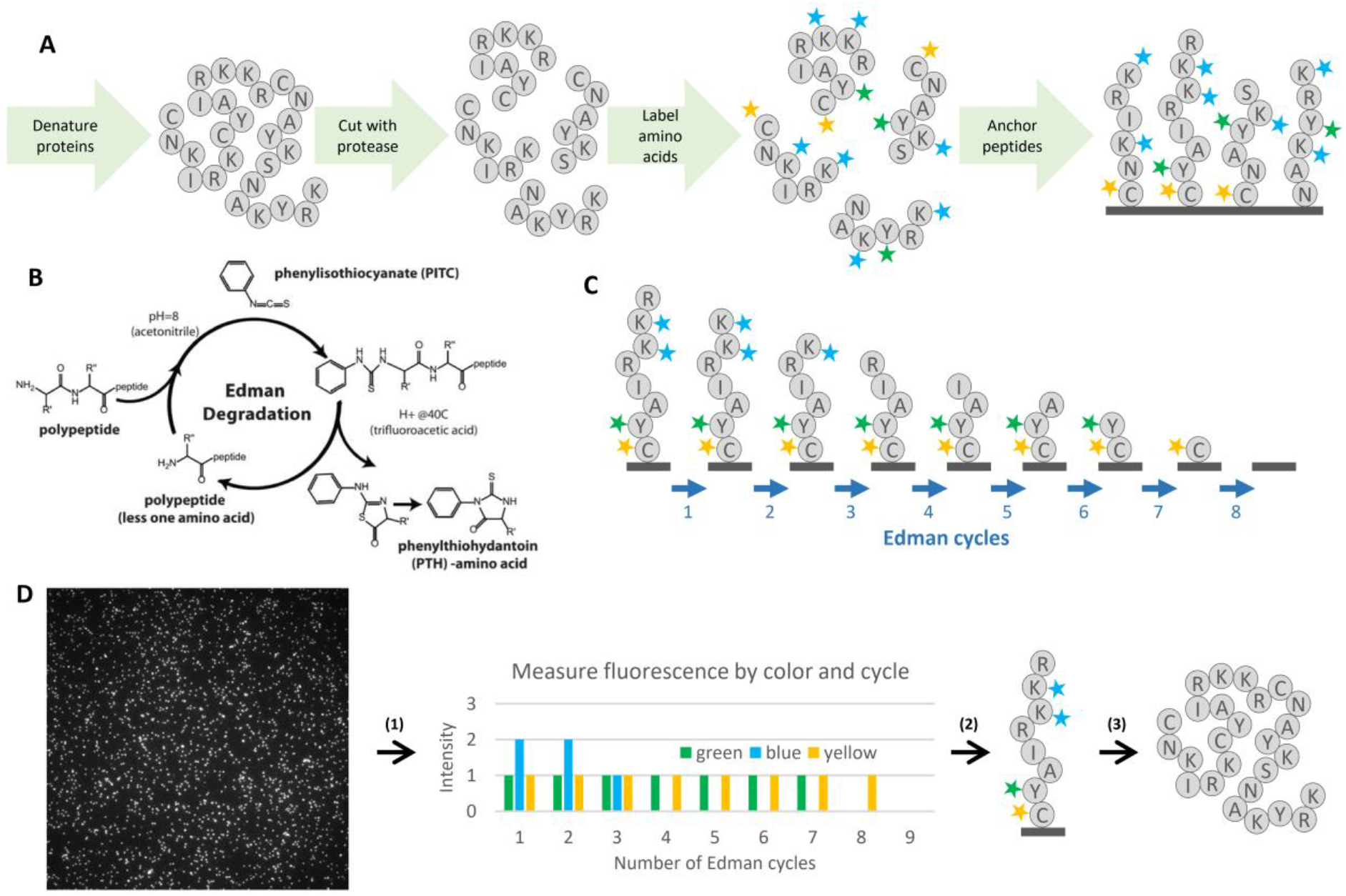
Overview of protein fluorosequencing. (**A)** illustration of the sample preparation process. Each grey circle represents an amino acid, and the letter in the circle corresponds to the standardized single letter amino acid codes. In the diagram, proteins are denatured, cleaved with protease, labeled with fluorescent dyes, and then labeled peptides are immobilized by their C-termini on the surface of a flow-cell. (**B**) The Edman degradation chemical reaction cycle, used to predictably remove one amino acid per cycle from each peptide. (**C)** For a given peptide, the sequencing process removes amino acids one at a time from the N-terminus, taking with them any attached fluorescent dyes. **D**: Major steps in computational data analysis include: **(1)** For each field of view, performing image analysis to extract fluorescence intensities for each spot (peptide) in each fluorescent channel across time steps (cycles), collating the fluorescence intensity data per spot across timesteps and colors. A vector of fluorescence intensities is produced, giving a floating-point value for every timestep and fluorophore color combination. **(2)** These raw sequencing intensity vectors (raw reads) must then be classified as particular peptides from a reference database. This step is the primary concern of this paper. **(3)**

In theory, this process gives a direct readout of each peptide’s amino acid sequence, at least for the subset of labeled amino acids (**Figure 1D**), but in practice there are several complications because of the single-molecule nature of this sequencing method. Single molecule fluorescence intensities are intrinsically noisy, arising from the repeated stochastic transitions of each individual dye molecule between ground state and excited state, making stoichiometric data inexact, particularly when there are large fluorophore counts. Typically, no more than 5-6 copies of the same amino acid, hence dye, are expected for average proteolytic peptide lengths, with the number of distinct colors (*i*.*e*., fluorescent channels) set by the microscopy optics and available dyes, here assumed to be 5 or fewer. However, inevitably with any chemical process, some fluorophore labeling reactions fail to occur, and photobleaching or chemical destruction can destroy fluorophores in the middle of a fluorosequencing run. At some low rate, peptides may detach from the flow-cell during sequencing, and Edman degradation can skip a cycle. These error rates, while individually small (approximately 5% each in published analyses [2]), collectively add difficulty to peptide identification, necessitating computational methods to process these data.

Currently, there are no published algorithms for mapping fluorosequencing reads to a reference proteome to identify the proteins in a sample. The first analyses of fluorosequencing data used Monte Carlo simulations to generate realistic simulated data as a guide for data interpretation and fitting of experimental error rates [1] [2]. While this strategy did not scale well computationally to full proteomes, it suggested that probabilistic modeling of the fluorosequencing process could provide a powerful strategy for interpreting these data. In this paper, we explore the application of machine learning to develop a classifier that correctly accounts for the characteristic fluorosequencing errors but is computationally efficient enough to scale to the full human proteome.

Viewing this as a machine learning problem is challenging due to the large numbers of possible peptides in many biological experiments. For example, in the human proteome, there are about 20,000 proteins, which when processed with an amino-acid specific protease such as trypsin can correspond to hundreds of thousands or even millions of distinct peptides, each of which can potentially vary due to post-translational modifications or experiment-specific processing. This puts fluorosequencing data analysis squarely in the realm of Extreme Classification problems, which are known to be challenging to handle in practice [9].

To analyze these data, we took advantage of the ability to generate simulated fluorosequencing data using Monte Carlo simulations [1] [2] to test k-Nearest Neighbors (kNN) classification and found it gave results of poor quality but is able to scale efficiently to the full human proteome while maintaining reasonable runtimes. These initial explorations motivated the developments presented in this manuscript, which focuses on the specific challenge of matching fluorosequencing reads to peptides from a reference proteome (*peptide-read matching*).

Here, we propose a specialized classifier which combines heavily optimized Hidden Markov Models (HMMs) to model the peptide chemical transformations during fluorosequencing, in combination with kNN pre-classification to reduce runtime. We call this tool *whatprot*, compare it with kNN and a classifier which uses HMMs without the kNN based runtime reduction, and demonstrate that the hybrid HMM-kNN approach offers a powerful and scalable approach for interpreting protein fluorosequencing data with the use of a reference proteome.

## Methods

### Monte Carlo simulation

To generate training and testing data typical of fluorosequencing experiments, we performed Monte Carlo simulations based on the model and parameters described in [1] [2]. These parameters are the dye loss rates *p*_*c*_, which differ for each color *c*, the missing fluorophore rates *m*_*c*_, the Edman cycle failure rate *e*, the peptide detachment rate *d*, the average fluorophore intensity *μ*_*c*_, and the standard deviation of fluorophore intensity *σ*_*c*_. We additionally model a background standard deviation 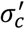. Based on prior estimates for the dye Atto647N ([1] [2]), we used the following values unless otherwise noted: *p*_*c*_ = .05, *m*_*c*_ = .07, *e* = .06, *d* = .05, *μ*_*c*_ = 1.0 (arbitrary rescaling of intensity values), *σ*_*c*_ = 0.16, 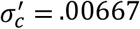. Although the code permits different values for different colors *c*, for our simulations, we modeled each color of fluorophore with identical error values for simplicity.

An overview of the process with definitions for key terms is provided in **Figure 2**. We generate simulated data in two formats. The first of these formats we refer to as a *dye track*, and it indicates the number of remaining fluorophores of each color at each time step after considering sequencing errors. Thus, each copy of one particular peptide sequence may give rise to a different specific dye track in a sequencing experiment depending on the details of the labeling schemes and sequencing efficiencies. To simulate a dye track, we randomly alter (with a pseudo random number generator) a representation of a dye sequence in a series of timesteps, writing to memory the count of each color of fluorophore as we progress until we reach a pre-set number of timesteps. In this simulation, we initially remove fluorophores with a probability of *m*_*c*_ before beginning sequencing. We then additionally perform a series of random events after logging fluorophore counts for each timestep: we remove the entire peptide and all fluorophores with a probability of *d* to simulate peptide detachment from the flow cell, we remove the last amino acid (and any attached fluorophore) with a probability of (1 − *e*), and we remove each fluorophore with a probability of *p*_*c*_, where *c* is the color of the fluorophore, to simulate fluorophore destruction. Each fluorophore count is stored as a two-byte numeric value.

**Figure 2.**
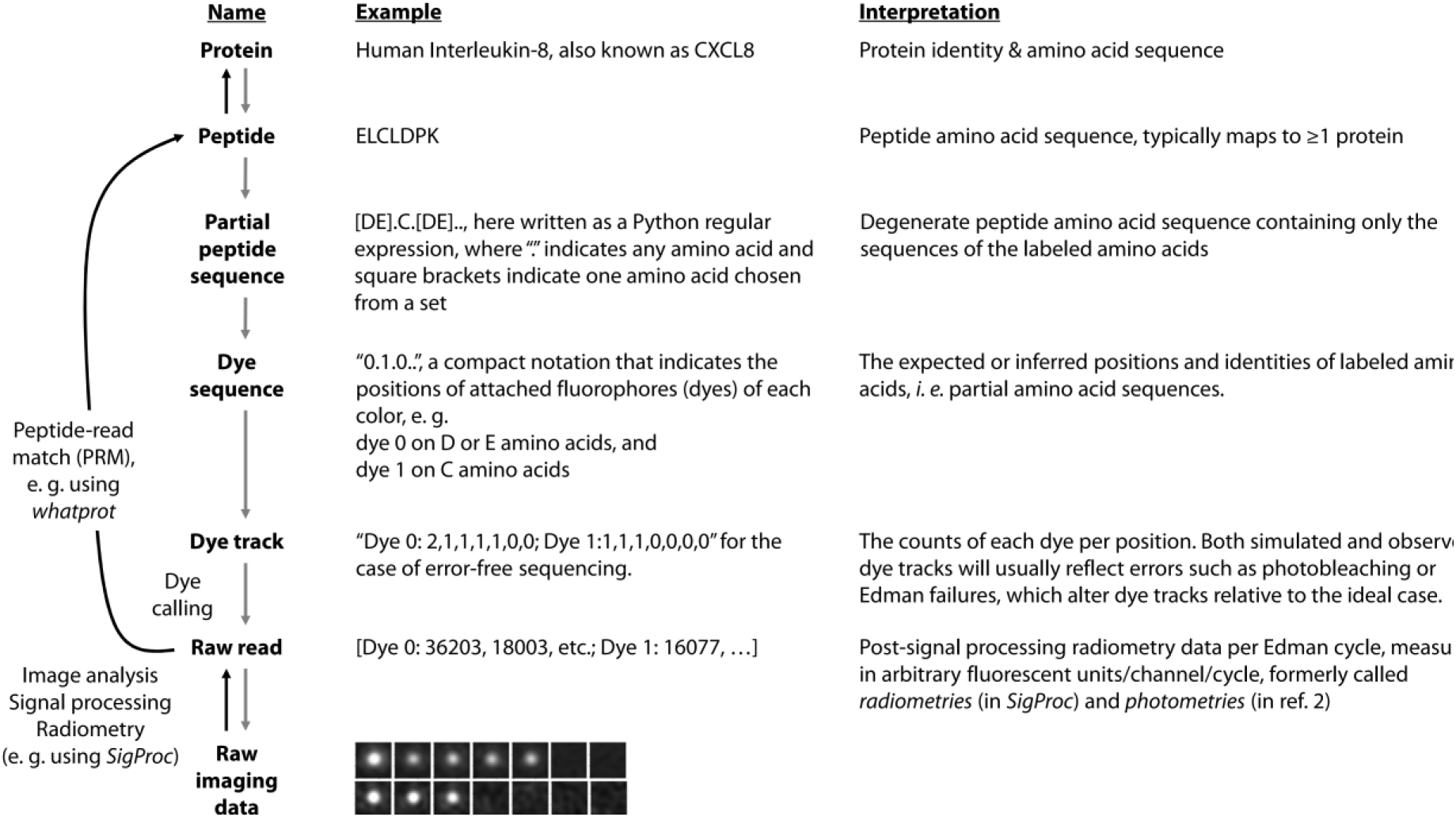
Nomenclature for different stages of fluorosequencing data analysis. The *whatprot* algorithm maps raw single-molecule protein sequencing reads to peptides and their parent proteins in the reference proteome (black arrows) by comparing experimental data (at bottom) to synthetic data generated using a Monte Carlo simulation (gray arrows).

The other format of data we consider is a *raw read*, which consists of radiometry data for each fluorescent color and Edman cycle. Raw reads result experimentally from signal processing and radiometry of the microscope imaging data from a fluorosequencing experiment. To simulate a raw read, we first simulate a dye track, and then we convert each fluorophore count into a double-precision floating point value indicating the fluorescent intensity. When we have a dye track entry indicating Λ_c_ fluorophores for a given fluorophore color *c*, we sample a normal distribution with a mean of Λ_*c*_*μ*_*c*_ and a variance of 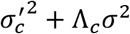. We perform this calculation for each channel at each time-step to simulate a raw read.

These radiometry *raw reads* simulate the fluorescent intensity data we would expect to collect from processing raw single molecule microscope images, a process currently performed for experimental data using the algorithm *SigProc* (Part of Erisyon’s tool *Plaster*, https://github.com/erisyon/plaster_v1), as in [10] [11].

### Bayesian classification with HMMs

Whatprot builds an independent HMM for each peptide in a provided reference proteome dataset. Each state in this HMM represents a potential condition of the peptide, including the number of successfully removed amino acids, and the combination of fluorophores which have not yet photobleached or been destroyed by the chemical processing (**Figure 3**). Transition probabilities between these states can be approximated using previously estimated success and failure rates of each step of protein fluorosequencing.

**Figure 3.**
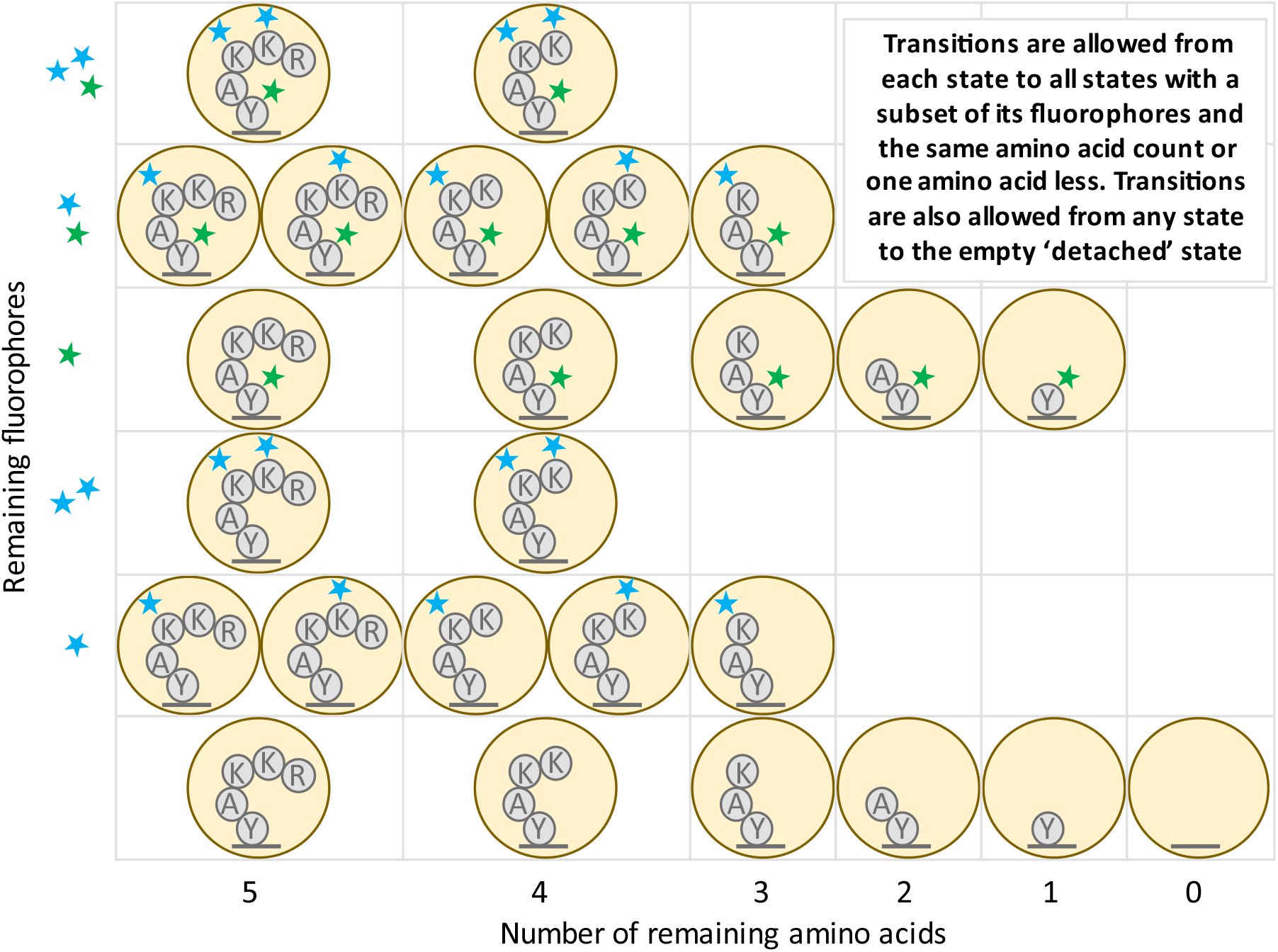
Illustration of the states and transitions of the HMM for an example peptide. For the amino acid sequence RKKAY, we illustrate the case where the lysine (K) residues are labeled with fluorescent dyes of one color (blue stars) and the tyrosine (Y) residue is labeled by a second color (green star).

We can use the HMM forward algorithm to associate a specific peptide to each *raw read* (a series of observed fluorescence intensities over time and across different fluorescence channels). We obtain the probability of the peptide given the raw read in two steps. First, we compute the HMM forward algorithm using each possible peptide in the dataset to obtain the probability of the raw read given each peptide. This uses the forward algorithm formula

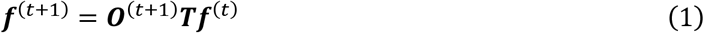

Where ***f***^(*t*)^ represents the cumulative probabilities for each state at timestep *t*, where 0 ≤ *t* ≤ *T*, ***O***^(*t*)^ represents the diagonal emission matrix for the observation seen at timestep *t*, and ***T*** represents the transition matrix which is the same at every timestep. The entries in each ***f***^(*t*)^, ***O***^(*t*)^, and in ***T***, represent the following probabilities:

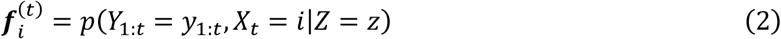

Where *Y*_1:*T*_ are the random variables for the observations, *y*_1:*T*_ are their true values, *X*_1:*T*_ are the random variables for the state in the HMM and *Z* is the random variable representing the peptide, and *z* is a value it can take. We also have diagonal matrices ***O***^(*t*)^ defined as:

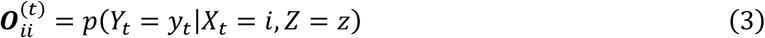

And :

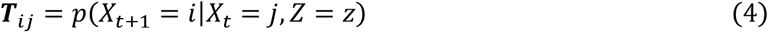

We start from an initial state ***f***^(0)^ which we compute by taking into account the missing fluorophore rate *m*_*c*_. Applying (1) repeatedly starting with the initial state ***f***^(0)^ yields a value for ***f***^(*T*)^, and we can sum the entries to compute:

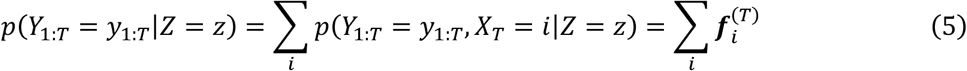

Then, by using Bayesian inversion to normalize the data, we compute the probability of the peptide given the raw read, as given by:

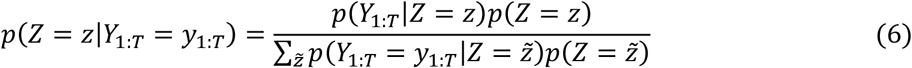

We implemented several algorithmic optimizations to this approach to reduce runtime. These included reducing the number of states in the HMMs, factorin the HMMs’ transition matrices into a product of matrices with higher sparsity, pruning the HMM forward algorithm to consider only reasonably likely states at each timestep, and combining the HMM classifier with a kNN pre-filter that can rapidly select a short-list of candidate peptides for re-scoring by the HMM. We implemented the linear algebra and tensor operations being performed in a manner that makes productive use of spatial and temporal locality of reference. We describe these optimizations in more detail in the following sections and in the supplemental **Appendices**.

### HMM state space reduction

We combine states of certain peptide conditions into more inclusive states in our model. For peptide conditions to be combined into these more inclusive states, they must have experienced the same number of *successful* Edman degradation events (so that they will have the same number of amino acids remaining), and they must have the same numbers of fluorophores of all colors. An example of the resulting HMM for a sample peptide is shown in **Figure 4**.

**Figure 4.**
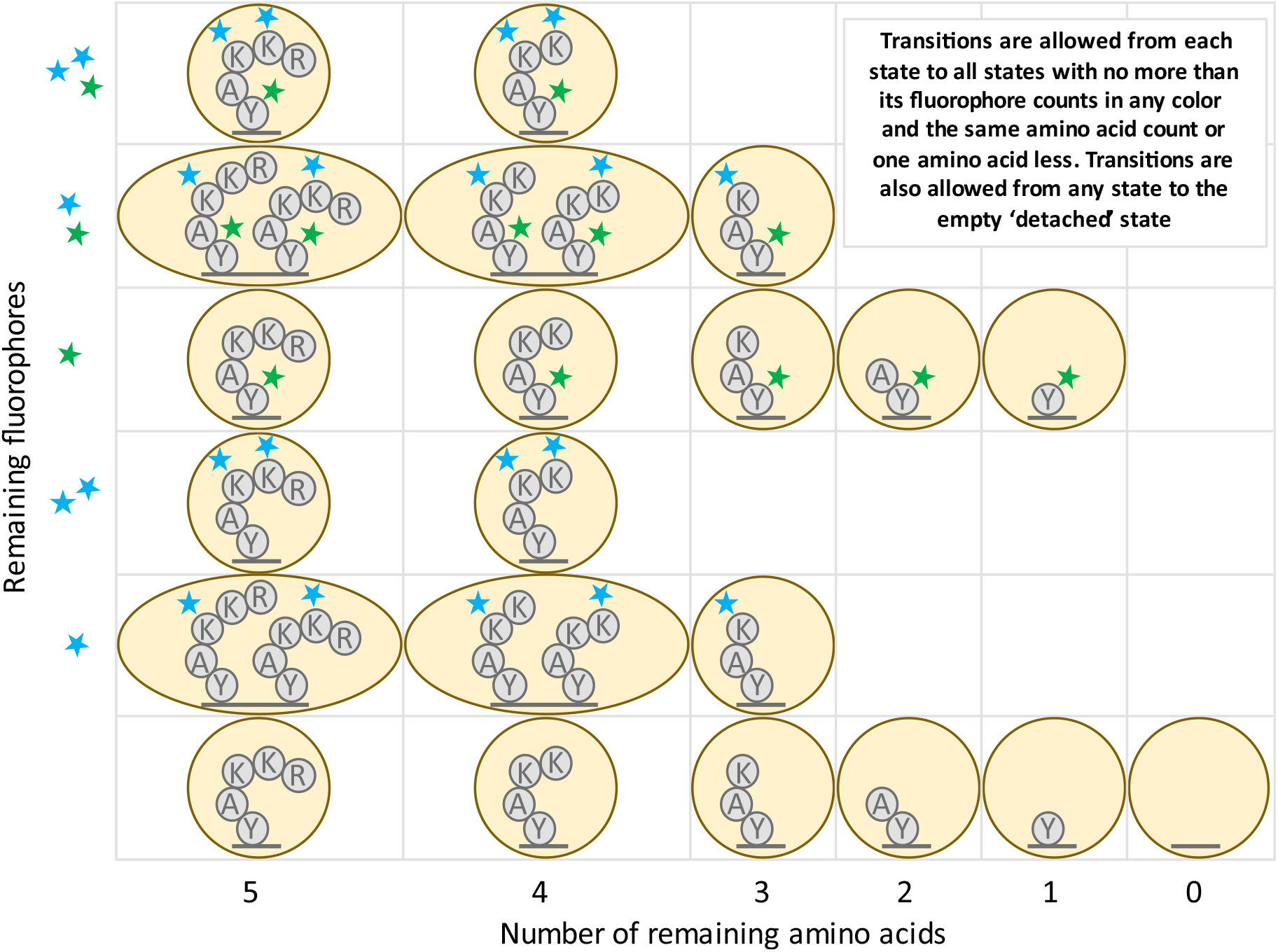
Illustration of HMM state space reduction for the peptide of Figure 3. States are combined that have both the same number of amino acids remaining and the same fluorophore counts for each color of fluorophore.

A similar state reduction to ours was previously described by Messina and colleagues in [12]. The reduction requires fluorophores to behave independently of each other, so that the status of one fluorophore is uncorrelated with the status of any other. While this is not true in practice due to FRET (Förster resonance energy transfer) and other dye-dye interactions, quantification of this effect in the imaging conditions used for fluorosequencing suggests that these effects are negligible enough to ignore [2]. The authors of [12] also require the fluorophores to be indistinguishable to reduce the numbers of states. This is not true in our case because we use Edman degradation and because we use multiple colors of fluorophores.

Nonetheless, we demonstrate in **Appendix A1** that despite these complications, this state space reduction incurs no loss in the theoretical accuracy of the model. Further, we demonstrate that this reduces the algorithmic complexity from what would otherwise be exponential with respect to the number of fluorophores, to instead be tied to the product of the counts of fluorophores of each color.

### Transition matrix factoring

In the HMM forward al orithm, a vector of pro a ilities with one value for each state in the HMM’s state space is repeatedly multiplied by a square transition matrix. This operation is the dominant contribution to the algorithmic complexity of the HMM forward algorithm. Therefore, by making multiplication by the transition matrix more algorithmically efficient, we can improve the theoretical complexity of our computational pipeline.

We factor this transition matrix into a product of highly sparse matrices. This factorization is done by creating a separate matrix for each independent effect under consideration, including loss of each color of dye (where each color is factored separately), Edman degradation, and finally, peptide detachment (**Figure 5**). As with the state space reduction, this optimization incurs no loss in the accuracy of the model, and furthermore, these matrix factors, even in combination, are far sparser than the original transition matrix when computed for larger peptides. This greater sparsity can be leveraged to achieve superior algorithmic complexity results (see **Appendix A2**).

**Figure 5.**
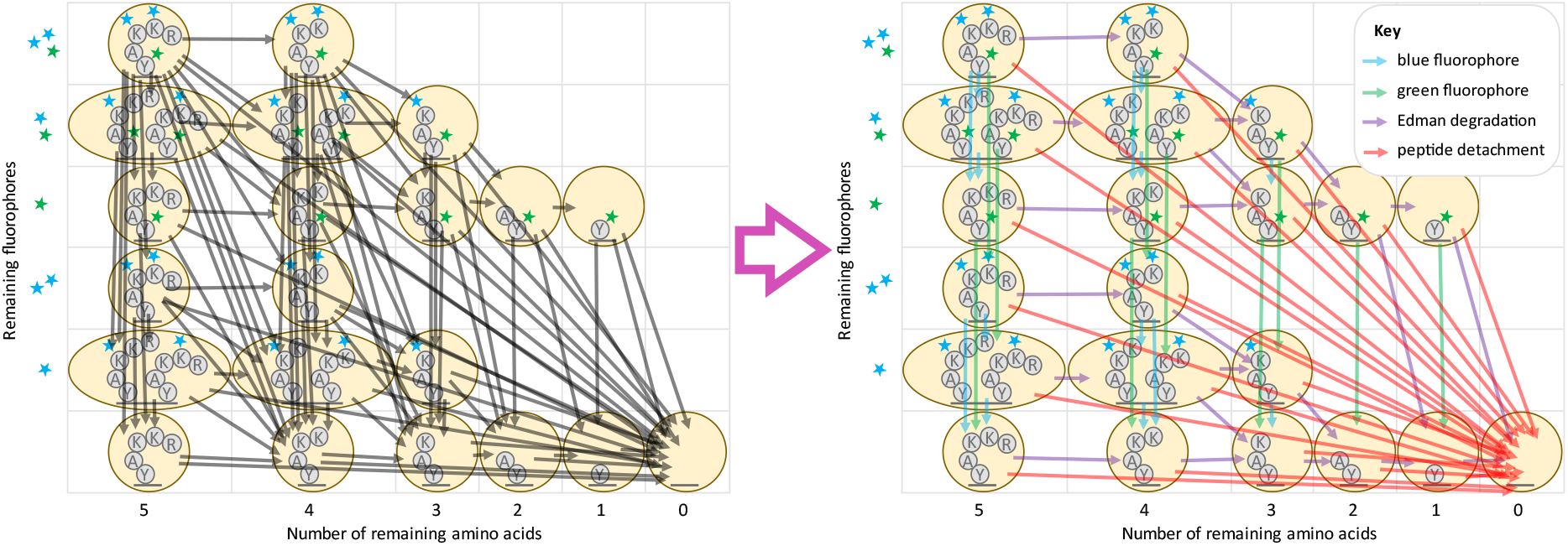
An illustration of the factoring of the transition matrix for the peptide from Figure 4. Note especially the reduction in the total number of transitions (arrows) when the transition matrix is factored. At left, black arrows represent non-zero entries in the unfactored transition matrix. At right, colored arrows (see key) represent non-zero entries in each of the matrices in the factored product. In both diagrams, arrows from a state to itself are omitted for visual clarity.

### HMM pruning

Despite significant improvements in the algorithmic complexity of an HMM for one peptide given so far from state space reduction and matrix factorization, performance can be improved if we consider approximations. Intermediate computations contain mostly values close to zero, which will have inconsequential impact on the result of the HMM forward algorithm. The most significant contribution to this occurs for the HMM emission calculations. While there may be many states of a peptide which have a significant probability of producing a particular observation value, in most states (particularly for larger peptides) the observed value is extremely unlikely.

Emission computations can be viewed as multiplication by a diagonal matrix, different for each emission in a raw read. The entries represent the probability of the indexed state producing the known emission value for that timestep. We prune this matrix by setting anything below a threshold to zero, which increases the sparsity of the matrix. Although use of a *naïve*, but standard, sparse matrix computational scheme would reduce runtime, we show that better algorithmic complexity can be achieved with a more complicated bi-directional approach in **Figure 6** and **Appendix A3**. While we did not implement this approach precisely, a consideration of this effect served as inspiration for a technique combining pruning with matrix factoring, as described next.

**Figure 6.**
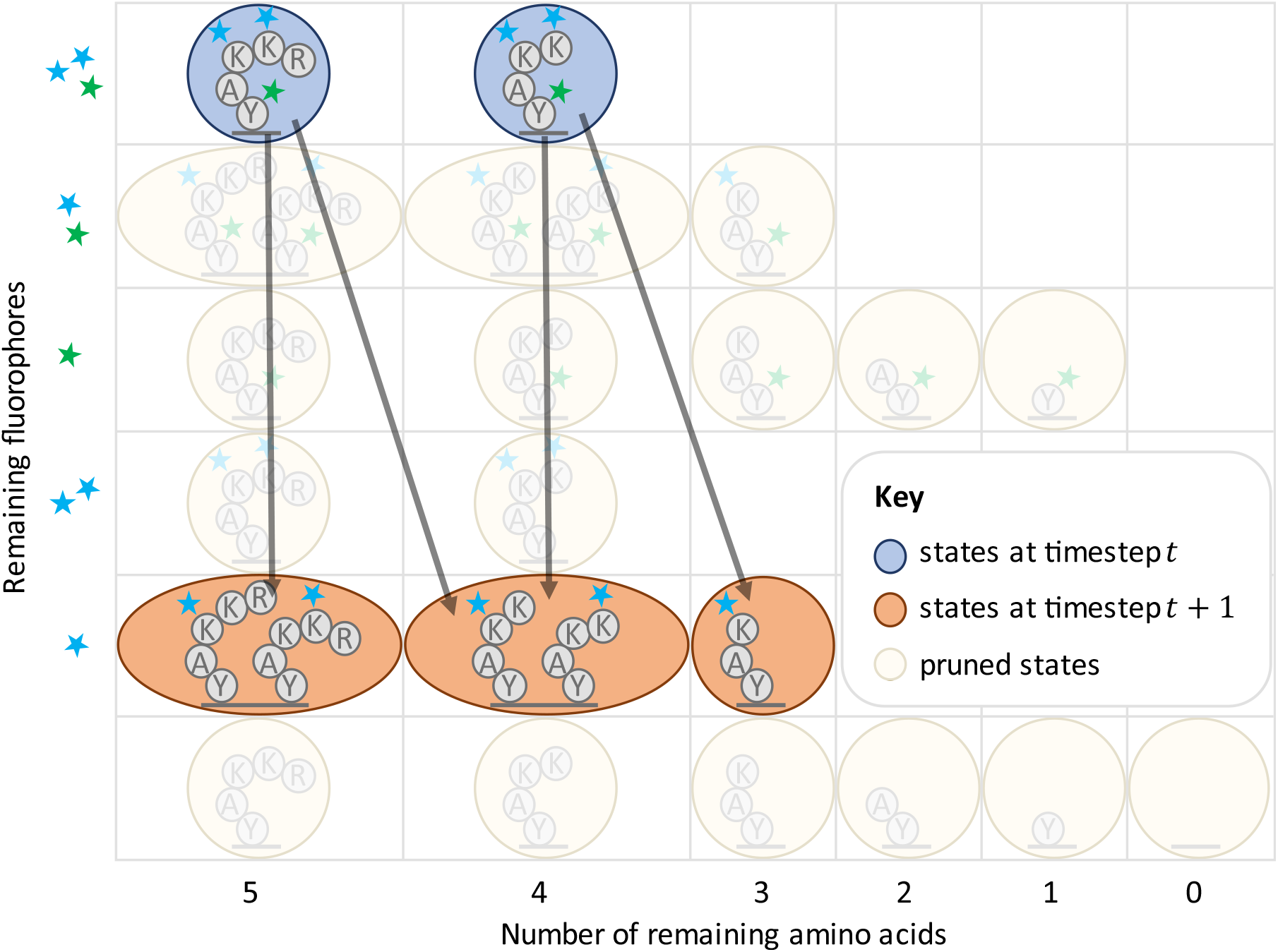
Illustration of the effects of HMM pruning for the peptide of Figure 5.

### Combining transition matrix factoring with HMM pruning

Both transition matrix factoring and HMM pruning appear, at first glance, to be incompatible improvements. These approaches can be combined, but the bi-directional sparse matrix computational scheme introduces significant additional difficulties.

We view the various factored matrices as tensors and propagate contiguous blocks of indices forwards and backwards before running the actual tensor operations to avoid unnecessary calculations (**Figure 7**). Contiguous blocks of indices are needed because propagating lists of indices across the various factors of the matrices has the same computational complexity as multiplying a vector by these matrices. This may make the pruning operation seem less optimal in a sense, as some values that get pruned may be bigger than some that are kept due to this form of indexing. Nevertheless, we found the tradeoff to be favorable in practice (**Appendix A4**).

**Figure 7.**
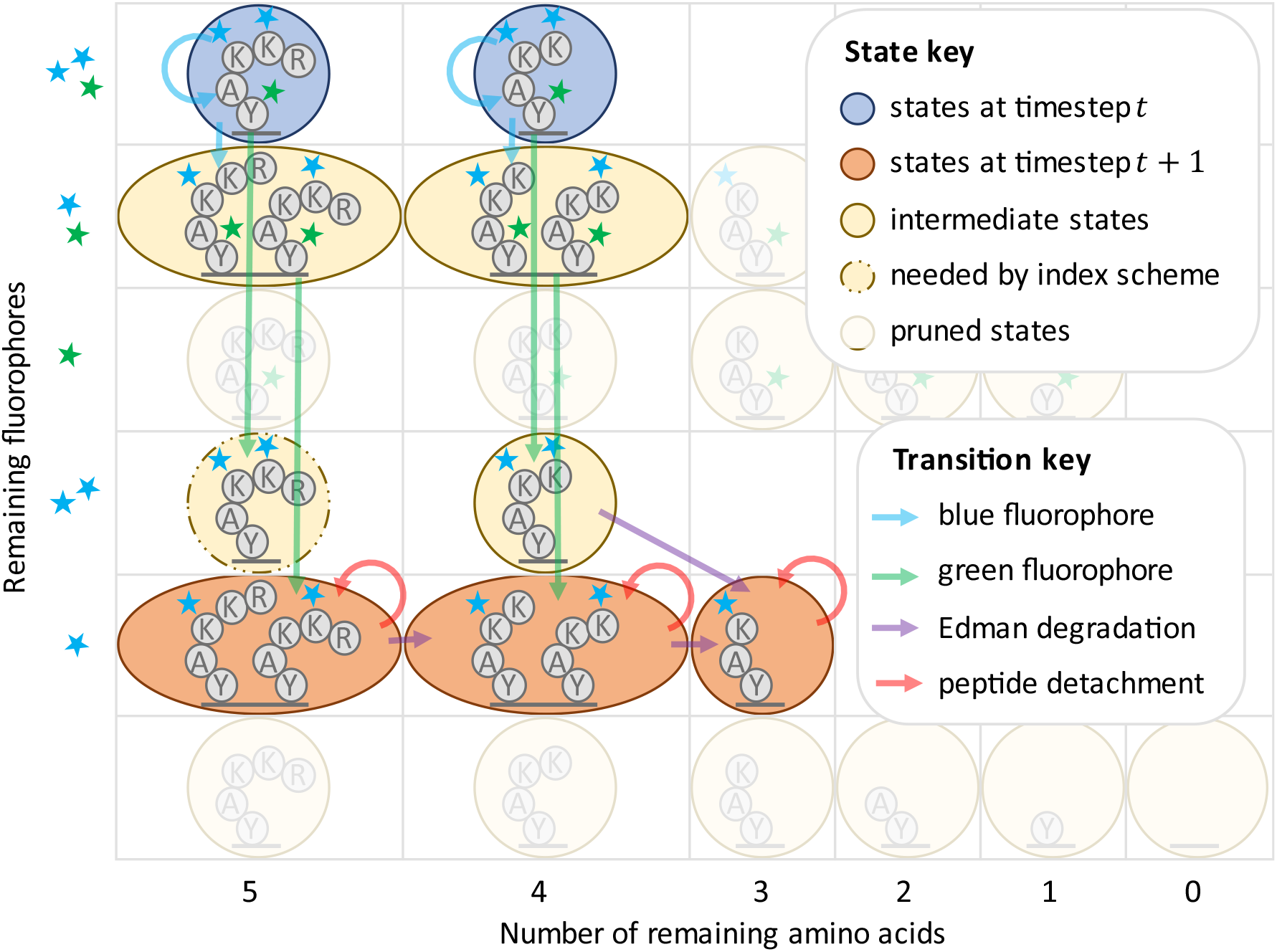
Illustration of HMM pruning combined with transition matrix factoring for the peptide of Figure 5. We emphasize that this is an anecdotal example; while there are more arrows here than in **Figure 6**, this strategy provides an improvement in asymptotic complexity, as described in **Appendix A4** and shown in experiments with simulated data.

Of note, instead of pruning by the raw values, we prune all states such that the known emission value is outside of their pre-configured confidence interval. In this way we provide some confidence that the fraction of true data inadvertently zeroed out is negligible.

### *k*-Nearest Neighbors classification

Most traditional machine learning classifiers have an algorithmic complexity which scales proportionally or worse to the number of classification categories. The Bayesian classifier we have so far described is no exception; each raw read must be compared against every peptide in the reference dataset to be classified. There are many problems in biology which require large reference datasets, human proteomic analysis being one example. The human proteome has 20,000 proteins, which when trypsinized generate hundreds of thousands of peptides. Classification against these many categories is computationally intractable with a fully Bayesian approach.

In contrast, the algorithmic complexity of kNN scales logarithmically with the number of training points used. For this reason, tree-based methods are common in other Extreme Classification applications [9], where similarly massive numbers of categories are under consideration. Unfortunately, the resulting faster runtimes come at a significant cost; kNN often gives far worse results in practice than a more rigorous Bayesian approach.

For purely kNN based classification, we simulate 1000 raw reads per peptide in the reference to create a training dataset and put these into a custom KD-Tree implementation for fast and easily parallelizable nearest neighbor lookups. We do not allow edits in our KD-Tree after it is built so as to allow parallelized lookups to occur without any concern for locks or other common issues in parallel data structures. We also reduce the memory footprint of the KD-Tree through an unusual compression scheme. For our training data, we use dye tracks instead of raw read radiometry data; this alone reduces the memory footprint of the KD-Tree by a factor of four (dye tracks have a two-byte numeric value for every timestep/color combination, while radiometry data have an eight-byte double-precision floating point number). But this allows another further compression technique; we find all identical dye tracks and merge them into one entry. With these dye tracks entries in the KD-Tree we store lists of peptides that produced the dye track when we simulated our training data, along with how many times each peptide produced that dye track.

To classify an unknown raw read, the *k* nearest dye track neighbors to a raw read query are retrieved. These neighbors then vote on a classification, with votes weighted using a Gaussian kernel function, 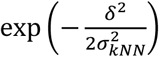, where *δ* is the Euclidean distance between the query raw read and the neighbor, and *σ*_*kNN*_ is a parameter of the algorithm. A neighbor is also weighted proportionally to the number of times it occurred as a simulation result and will split its voting weight among all of the peptides that produced that dye track proportionally to the numbers of times each peptide produced the dye track during simulation of training data.

Once voting is complete, the highest weighted peptide is then selected as the classification, with its classification score given as a fraction of its raw score over the total of all the raw scores. We have explored multiple choices of *k* and *σ* values to optimize the performance.

### Hybridizing kNN with Bayesian HMM classification

To combine the computational efficiency of kNN with the accuracy of the HMM model, we defined a classifier which hybridizes these two disparate methods. We use a kNN classifier to reduce the reference dataset, for each raw read, down to a smaller shortlist of candidate peptides. These candidates can then be used in the Bayesian classifier by building HMMs to compare them against the specific raw read.While this can result in the true most likely peptide not being in the shortlist and therefore not being selected by this hybrid classifier, with a sufficiently long shortlist this is highly unlikely. A larger problem is in performin Bayes’ rule, as in (6). n e act formula for Bayes’ rule requires an e haustive set of probability values for every potential outcome, which are summed in the denominator. Avoiding determining every probability makes this impossible. Instead, we can estimate Bayes’ rule as follows:

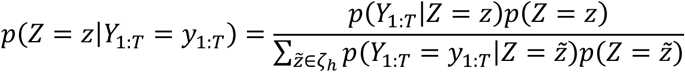

Where *ζ*_*h*_ is the set of up to *h* peptides selected by the kNN method; we require *z* ∈ *ζ*_*h*_. Although we lose theoretical guarantees of optimal accuracy given the model, this change provides a considerable improvement to the algorithmic complexity. The algorithmic complexity to classify one raw read using a fully Bayesian approach is *O*(*RW*), where *R* is the number of peptides in the reference dataset, and *W* is the average amount of work needed to run an HMM for one peptide fluorophore combination. In comparison, with the hybridized classifier, the algorithmic complexity is *O*(log(*RQ*) + *hW*) where *Q* is the number of raw reads in the training dataset simulated for each possible peptide.

We chose specific values for *h, σ*, and *k*, by comparing the runtime and PR curves on simulated datasets.

### Maintaining spatial locality of reference

Spatial and temporal locality of reference is the tendency of some computer programs to access nearby data points at similar times. Modern CPUs are designed to make this extremely efficient through multi-level batch caching schemes which cache data from RAM that is nearby a memory address being accessed, so that nearby data can be read more quickly. Programs which exploit this in read-write intensive pieces of code can often achieve significant runtime acceleration compared to programs which do not.

We wrote highly optimized kernel functions to perform our structured matrix/tensor operations which exploited the sparse nature of the problem while also iterating over elements in what we believe to be an optimal or near optimal fashion for most computer architectures. We believe this provided considerable improvements in performance, though this has not been rigorously tested.

## Results

We simulated the fluorosequencing of peptides to obtain labeled training and testing data (**Figure 2**). We generated several datasets, each with a randomized subset of the proteins in the human proteome. We selected 20 proteins (.1% of the human proteome), 206 proteins (1% of human proteome), 2,065 proteins (10%), and 20,659 proteins (the full human proteome). We repeated this randomized selection scheme to examine several protease and labeling schemes. These were (1) trypsin (which cleaves after lysine (K) and arginine (R) amino acids) with fluorescent labels for aspartate (D) and glutamate (E) (these share a fluorophore color due to their equivalent reactivities), cysteine (C), and tyrosine (Y), (2) cyanogen bromide (which cleaves after methionine (M) amino acids) with D/E, C, Y, and K, (3) EndoPRO protease (which cleaves after alanine (A) and proline (P) amino acids) with D/E, C, and Y, (4) EndoPRO with D/E, C, Y, and K, (5) EndoPRO with D/E, C, Y, K, and histidine (H). Thus, in the schemes examined, of the 20 canonical amino acid types found in most proteins, either one or two were recognized by the protease and up to 6 additional amino acids were labeled by fluorescent dyes.

These databases of peptides were used to generate databases of idealized *dye sequences, dye tracks*, and *raw reads*, used for training and testing purposes for the various models. For our test data for each dataset, we generated 10,000 raw sequencing reads by randomly selecting peptides with replacement from the dataset and simulating sequencing on them using the methods described in the *Monte Carlo simulation* section of this paper. For both dye tracks and simulated fluorescent intensity measurements, results where there were zero fluorophores throughout sequencing were discarded, as these would fail to be observed in an actual sequencing experiment.

We collected and compared runtime data and precision-recall curves for several different purposes. With the trypsinized 3-color dataset, we performed a parameter sweep of the pruning cut-off for the HMM Bayesian classifier (**Figure B1**). Losses in precision and recall performance were negligible for cut-off values of 5 and greater, though the precision recall curves grew worse at smaller values. Runtimes shrank rapidly as the cut-offs were decreased. The pruning cut-off parameter sweep was also performed on the 20-protein cyanogen bromide dataset (**Figure B2**). We saw that in this second dataset, runtime improvements for lower cut-offs were even more extreme; a speed-up factor of about 1000 could be achieved with minimal effects on the precision recall plots. From these two simulations, we chose a cutoff value of 5 as providing the optimal trade-off between runtime and precision recall performance.

On the trypsinized dataset (full human proteome), we also swept the *k* and *σ*_*kNN*_ parameters of the NN classifier (**Figures B3, B4**). Here large values of *k* introduce modest reductions in precision recall performance, while the model is extremely sensitive to the selection of *σ*_*kNN*_. Based on this analysis, we suggest that good choices of these parameters are *k* = 10 and *σ*_*kNN*_ = 0.5.

We swept all parameters of the hybrid classifier for the trypsinized dataset as well (hybrid *h* parameter, *k, σ*_*kNN*_, and cut-off) (**Figures B5-8**). Here, the HMM cut-off parameter had less impact on runtime than for the pure HMM Bayesian classifier, but we still found a cut-off of 5 to be optimal. Higher values of *k* improved precision recall performance for the hybrid model, contrary to the results of the NN classifier on its own, and we therefore suggest setting *k* to 10000. *σ*_*kNN*_ had minimal impact of any kind, in contrast to its significant impact on the precision recall of the NN classifier; we nevertheless chose to set it to 0.5 in light of the data from parameter tuning for the NN classifier on its own. We also found that higher values of *h* improved performance, though the impact plateaus after *h* of about 1000, and we used that value for later experiments.

After tuning parameters, we compared the performance of the different classifiers when applied to 10,000 simulated fluorosequencing reads of peptides drawn randomly from all tryptic peptides in the human proteome (**Figure 8**). The hybrid classifier achieved similar precision recall curves to the Bayesian HMM classifier, which was much better than the precision recall curve of the NN classifier. The hybrid classifier also achieved runtimes of the same order of magnitude as the NN classifier, which was significantly faster than the runtime of the Bayesian HMM classifier.

**Figure 8.**
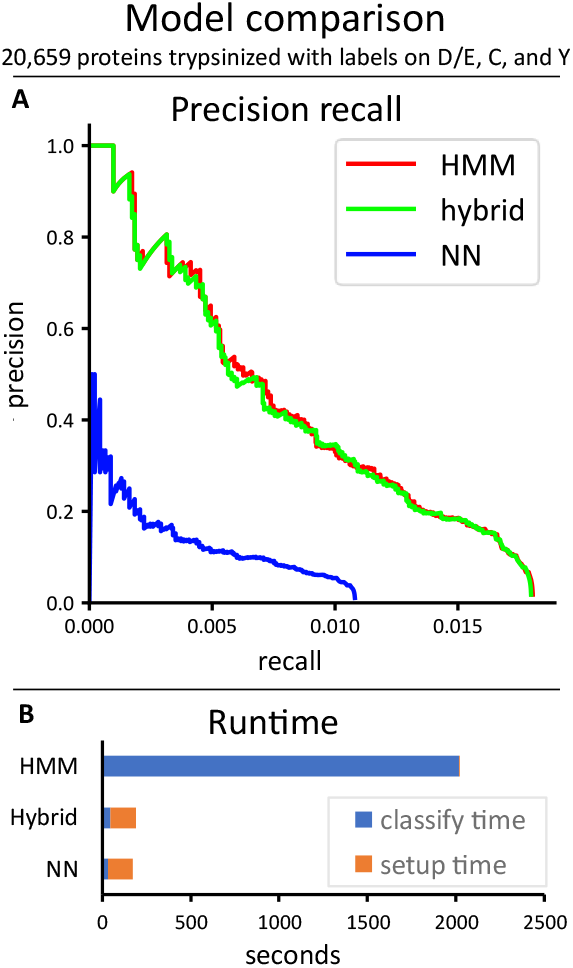
Comparison of the HMM (Bayesian), hybrid, and NN classifiers on a dataset of 10K reads from peptides chosen randomly from all 20,659 human proteins trypsiniz3ed38 and labeled on D/E, C, and Y. (**A)** The Precision recall curves. (**B)** Runtimes.

We also studied how the number of fluorophore colors affected the runtime and precision/recall of the hybrid classifier (**Figure 9**). We found that improvements in precision recall were possible with each additional color of fluorophore, but this did come at the cost of longer runtimes.

**Figure 9.**
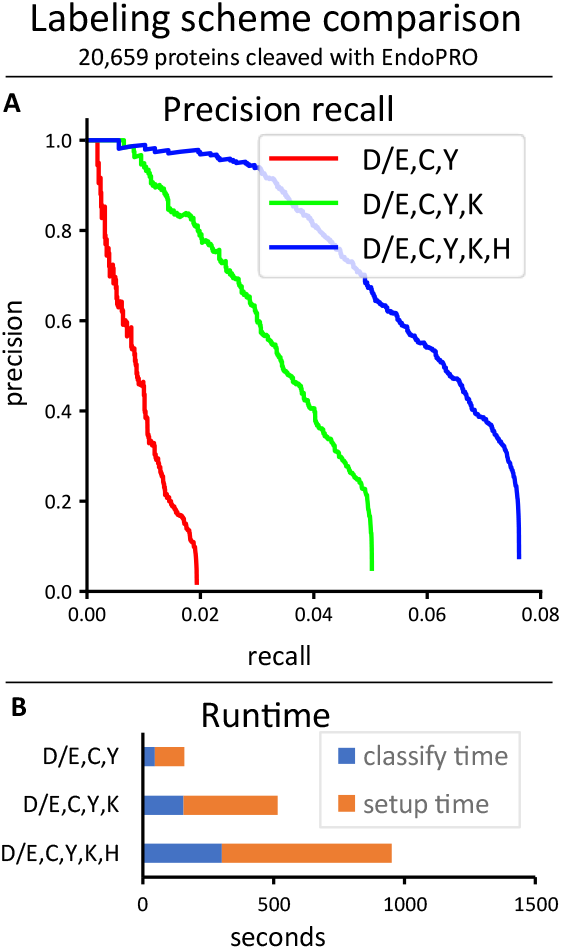
Comparison of the hybrid classifier on a dataset of 10K reads from peptides chosen randomly from all 20,659 human proteins cleaved with EndoPRO and labeled with three different labeling strategies. (**A)** The precision recall curves. (**B)** Runtimes.

We also investigated the effect of varying sizes of reference proteomes on the hy rid classifier’s performance, using the three color trypsinized dataset (**Figure 10**). We found that significantly better performance was possible when the reference database was smaller.

**Figure 10.**
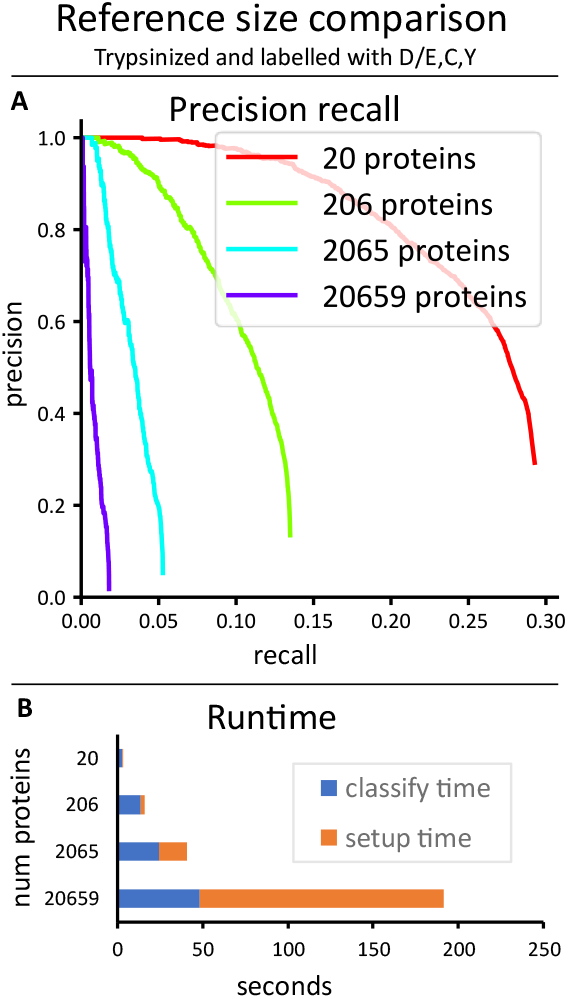
Comparison of the hybrid classifier across datasets of 10K reads each from peptides chosen randomly from different numbers of proteins treated with the same protease and labeling scheme. (**A)** The precision recall curves. (**B)** Runtimes.

The precision/recall curves plotted above (**Figures 8-10**) show the actual precision/recall scores based on data with known peptide classifications. When working with real data this will typically not be possible, because the real classifications will not be known. It is therefore important that the assignment probabilities produced by the classifier be well-calibrated, so that an estimate of the precision/recall (or as is more often the case in protein mass spectrometry, the false discovery rate (FDR)) can be computed in the absence of known labels. We verified that the probabilities output by the hybrid HMM classifier were indeed well-calibrated relative to the true assignment probabilities (**Figure 11A**). This in turn allowed us to compute a predicted precision/recall curve assuming that each classification is fractionally correct with a probability given by its classification score. A comparison of this predicted P/R with the actual precision/recall curve for the same set of reads shows excellent agreement (**Figure 11B**).

**Figure 11.**
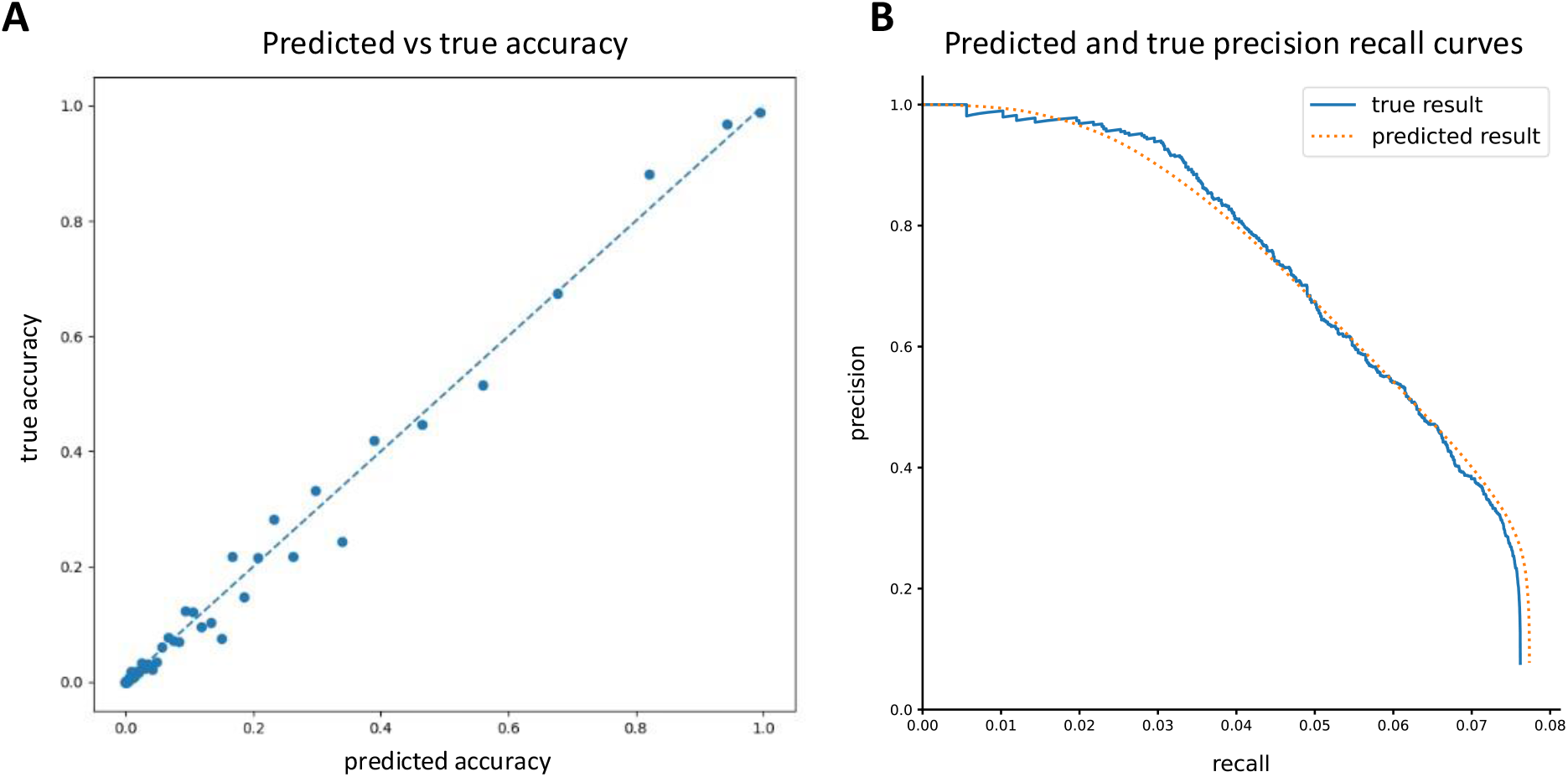
Analysis of the accuracy of probability estimates given as scores by the classifier. Based on 10K reads from peptides chosen randomly from all 20,659 human proteins cleaved with EndoPRO and labeled on D/E,C,Y,K,H. **(A)** Classification results were sorted by their predicted accuracy scores, and then equally distributed between 100 buckets. The average predicted and true accuracy scores were then computed for each bucket and plotted. **(B)** The true result precision/recall curve was computed as normal, while the predicted result precision/recall curve was plotted assuming each classification was fractionally correct according to its predicted accuracy score.

Whatprot specifically attempts to assign each raw fluorosequencing read to one or more peptides from the reference database, *i*.*e*., to identify and score *peptide-read matches* (PRMs), a process highly analogous to analytical interpretation of shotgun mass spectrometry (MS) proteomics data in which a key step is comparing experimental peptide mass spectra to a reference proteome (finding *peptide-spectral matches*, or PSMs [13]). However, observing multiple reads mapping to the same peptide will tend to increase the confidence that peptide is present in the sample, just as observing multiple peptides from the same protein will similarly increase confidence in that protein being present. Thus, we asked if considering the PRMs collectively led to performance increases for identifying peptides and proteins.

As shown in (**Figure 12**), proteins can be identified correctly at much higher rates than peptides, which are similarly identified at higher rates than individual reads. In fact, provided that a protein possesses some well-identified peptides that are unique, it can typically be identified with very high accuracy. For this test, we used a very simple protein inference scheme. First each peptide was scored to the maximum score of all reads identifying it, while penalizing reads which identified more than one peptide (dividing by *n* if *n* peptides were identified). Second each protein was scored as the maximum score of all peptides it contains, penalizing peptides which are associated with more than one protein (again dividing by *n* if *n* proteins were associated). However, the problem of integrating peptide level observations to protein observations has been studied extensively for MS [14] [15], and it is likely that these techniques will offer similarly strong interpretive power to the case of single molecule protein sequencing.

**Figure 12.**
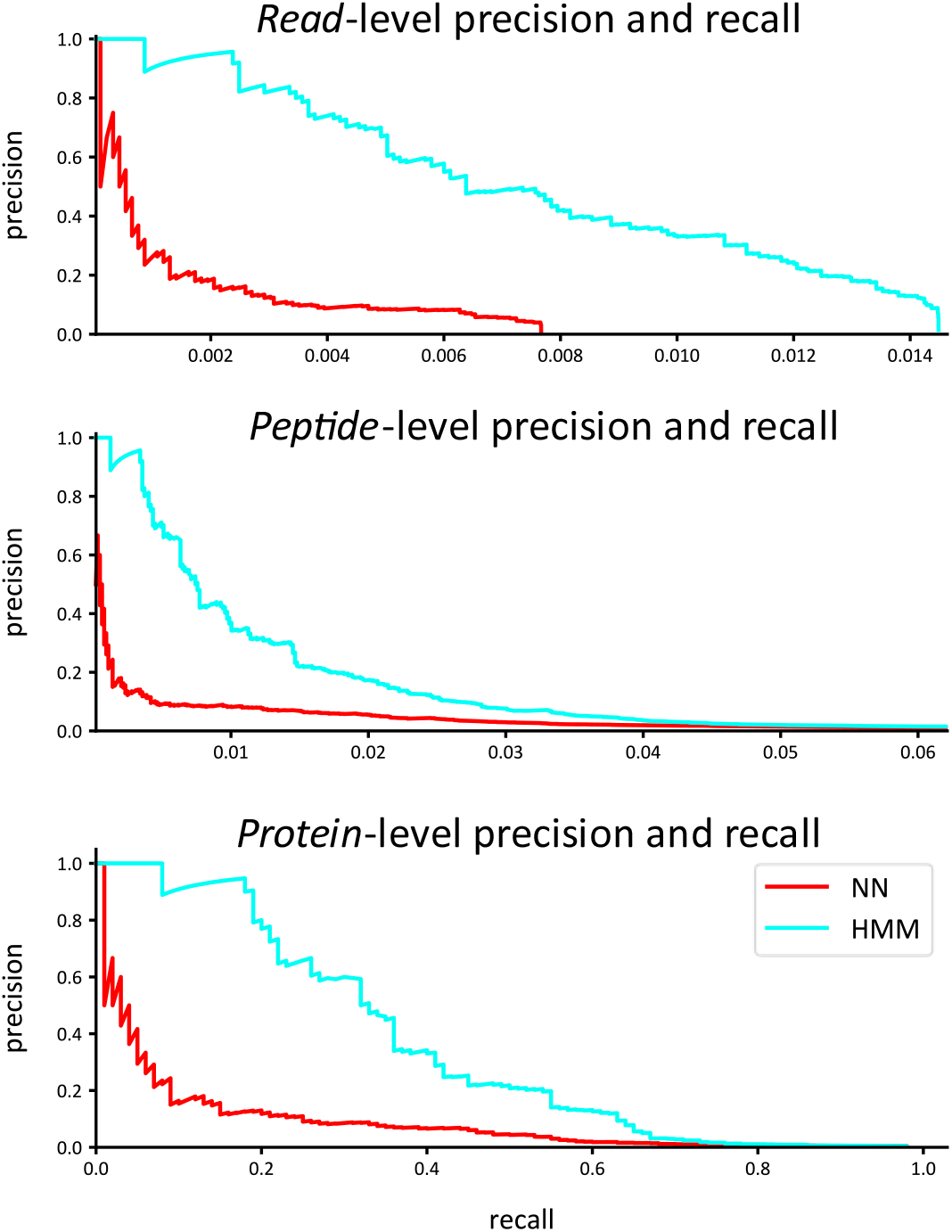
Precision and recall are improved for proteins by integrating identifications across peptides. The example shows 10K reads from peptides derived from 100 proteins randomly selected from the human proteome, considering trypsin digestion and labels on D/E, C, and Y.

## Discussion

We developed an HMM for interpreting single molecule protein fluorosequencing data and showed that a hybrid HMM/kNN model can achieve a high precision and recall comparable the HMM alone while maintaining a runtime comparable to the much faster kNN.

It is worth emphasizing that these analyses were performed on datasets of 10,000 raw fluorosequencing reads. In practice, users will likely want to analyze millions to billions of reads, so runs that completed in a seemingly reasonable amount of time might still be intractable in these scenarios with larger datasets, or at a minimum require computing clusters with high parallelization. For analyzing the current datasets in the runtime charts (**Figures 8-10**), note that the *blue* part of the bar graphs indicates the classify time, which will scale with the number of reads being classified (if all else remains equal), and the *orange* part of the graph indicates setup time, which should remain constant regardless of the number of reads (though it changes depending on the model and the size of the reference set).

It is also interesting that the HMM pruning operation is more necessary with longer peptides and more colors of fluorophores; with the trypsinized dataset labeling D/E, C, and Y, omitting pruning had little consequence, but in moving to cyanogen bromide with D/E, C, Y, and K, we observed a runtime speedup of about 1000-fold.

Finally, our data demonstrates that with a proper selection of parameter values, the hybrid model can achieve precision and recall performance virtually identical to the HMM Bayesian approach alone, while providing those results in a fraction of the time. Similarly, the pruning operation employed in the HMMs has no noticeable positive or negative effect on the precision recall curves while providing a considerable improvement in runtime performance.

A number of analytical techniques common in related fields were not explored. In tandem mass spectrometry (MS/MS), peptide spectral mapping is typically done either through database lookups and/or the use of simulated outcomes. Simulated mass spectra, consisting of ion pairs for the C- and N-terminal fragments for each potential breakage point, can be compared to the real data collected from the instrument [13]. Recent advances have been achieved by using deep learning to predict fragmentation behavior with higher quality than is possible with more traditional methods [16][24].While we use the notion of matching fluorosequencing reads to a reference database, the specific algorithms are distinct.

Nevertheless, of possible relevance from the field of MS/MS is the analysis of the false discovery rate (FDR) [25][26]. The FDR is affected by two distinct sources: a peptide may be misattributed to the wrong peptide, even when the true peptide is present in the reference dataset, and MS/MS datasets contain significant amounts of modified peptides or contaminants, whose spectra may be mistakenly assigned to peptides in the reference set [27]. FDR is typically evaluated using a decoy database, such as is generated using reversed proteins from the target database. The FDR can then be set by referring to the number of hits in the decoy database given a particular score, as the decoy database is designed such that it should in theory never have hits for the biological sample being analyzed [17]. While an estimate of FDR based in theoretical analysis of the problem could find the misattribution rate of true peptides, even this estimate would be incomplete, because there are errors in mass spectra of peptides that cannot be accounted for by existing theory; furthermore, any effect of modifications or contaminants would likely be omitted.

The utility of a similar decoy database strategy for estimating FDR for fluorosequencing is unknown and remains to be established. We note however that, due to the rigorous probabilistic nature of our analysis, a reasonable estimate of FDR can be performed by subtracting the sum of PRM scores from the number of PRMs. This is the same as one minus the precision in a predicted precision/recall curve, and the proximity of our predicted precision/recall curve to the real curve for a known dataset demonstrates the feasibility of this approach (**Figure 11**). This analysis likely fails to account for the contributions of modifications and contaminants. We therefore plan to explore this problem more extensively in future work.

We also considered techniques for DNA sequence reconstruction. In general, DNA sequencing provides *de novo* sequence reconstructions and does not use reference database matching, and therefore is not a good model for fluorosequencing. Nevertheless, base calling strategies may have some relevance. For example, methods for base-calling from conventional (e.g. Illumina style) DNA sequencing are straightforward [18] [19], and although errors occur, they are rare [20]. Analysis of errors in DNA sequencing is typically performed using multiple sequence alignment or k-mer based methods [21].Because the error rates are typically much lower in DNA sequencing than in fluorosequencing, we believe existing software is unlikely to be effective in this new domain.

Nanopore DNA sequence analysis methods could also be considered. Nanopores, similar to fluorosequencing, deal with single molecule data and the concomitant statistical noise that process involves. However, nanopore data is on a real time continuum, with a DNA fragment which may move through the nanopore at variable rates during sequencing. Base-calling, the assignment of nucleic acid bases to chunks of sequencing information, is again the step most analogous to fluorosequencing. State of the art base-calling methods for nanopore sequencing typically use either HMMs or recurrent neural networks (RNNs). Comparisons of existing approaches suggest that RNNs slightly outperform HMMs in this domain [22]. While this suggests that RNNs are worth exploring for fluorosequencing data, we have avoided this approach for two reasons. First, RNNs are a deep learning technique, which invariably requires access to massive amounts of data; this is not currently feasible with fluorosequencing unless that data is simulated. Second, our approach suggests the possibility of direct estimation of parameters using some variant of the Baum-Welch algorithm adapted to our use case, which we believe would be significantly more difficult in an RNN based approach [23].

## Conclusions

We have developed a powerful computational tool for the analysis of protein fluorosequencing data, which significantly increases the complexity of applications available to this new technology. This tool includes critically important optimizations which make our approach feasible in practice.

In future work we plan to implement a variation of the Baum-Welch algorithm to fit the parameters to data from a known peptide. We also wish to explore peptide and protein inference methods using peptide data classified using these methods. We may also explore *de novo* recognition of labels without use of a reference database.

## Acknowledgements

Code is available at https://github.com/marcottelab/whatprot. The authors gratefully acknowledge Jagannath Swaminathan, Angela Bardo, Brendan Floyd, Daniel Weaver, and Eric Anslyn for helpful guidance and discussion throughout the course of this project. M.B.S. acknowledges support from a Computational Sciences, Engineering, and Mathematics graduate program fellowship. E.M.M. acknowledges support from Erisyon, Inc., the National Institute of General Medical Sciences (R35GM122480), the National Institute of Child Health and Human Development (HD085901), and the Welch Foundation (F-1515).

## Appendix A

**A – detailed descriptions and proofs of algorithms**

### A0 – crib sheet for variables used throughout appendix A

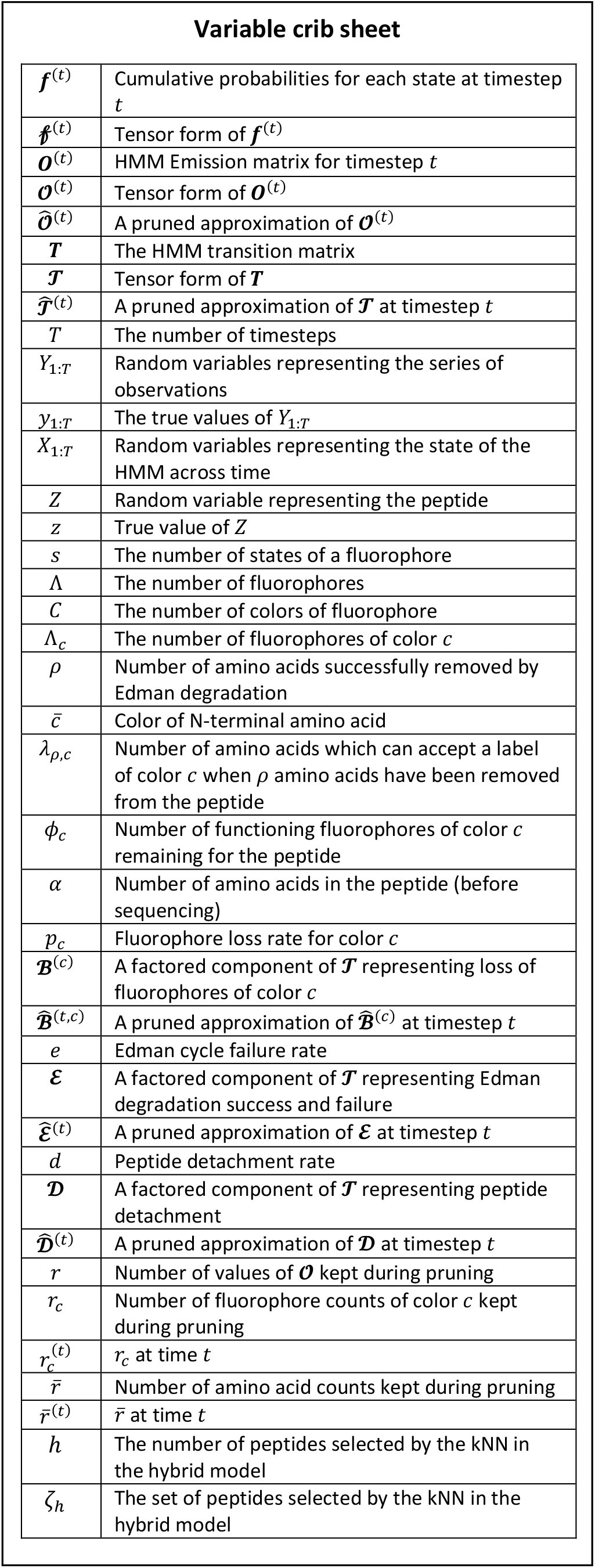

#### A1 – HMM state space reduction

#### The basics

For fluorophores that are distinguishable and dependent on each other, where *s* is the number of states of one fluorophore, and Λ is the number of fluorophores, the number of states of the whole system is given as in [12] by:

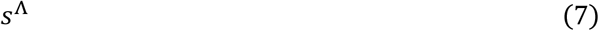

In contrast, if the fluorophores are indistinguishable and independent of each other, the number of states in the system is instead given by the combinatoric equation from [12]:

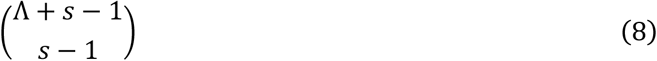

Imaging is typically performed with high concentrations of the antioxidant Trolox [2] and for relatively short time intervals (100 msec); reversibly photobleached fluorophores do not occur frequently in our data and we ignore them as a first approximation. Therefore, we take *s* to be 2, with one state for a functioning fluorophore, and another for a missing, photobleached, or chemically destroyed fluorophore. This reduces (8) to Λ + 1 states given Λ indistinguishable and independent fluorophores.

#### More colors

Obviously, a red fluorophore is distinguishable from a blue one. But we would still like to benefit from the indistinguishability of red fluorophores from red fluorophores, and of blue fluorophores from blue fluorophores. This is modeled by first considering the states for each color of fluorophore independently, and then taking the cartesian product of these state spaces. For *C* colors of fluorophore, where Λ_*c*_ is the number of fluorophores of color *c*, this results in the number of states given by:

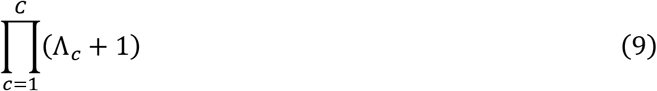

Each state then represents the number of remaining active fluorophores for each of our *C* colors of fluorophore.

#### Edman degradation

Our second challenge is the inclusion of Edman degradation in the HMM. Sequential removal of the N-terminal amino acid from each peptide breaks the assumption of indistinguishable fluorophores, which is the basis for the state reduction performed in [12]. However, through inductive reasoning we show that our model meets a weaker criterion, which can be used to merge states together as desired:

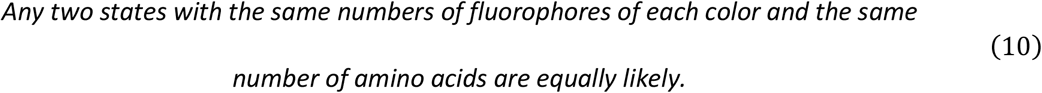

If we ignore Edman degradation, this follows directly from the assumed indistinguishability property of fluorophores of the same color; if two fluorophores behave identically, they are equally likely to be missing, photobleached, or chemically destroyed, thus it follows by symmetry that any two states with the same numbers of indistinguishable fluorophores of each color are equally likely. If we consider Edman degradation, then (10) is true for all states where no amino acids have yet been successfully removed. Let *ρ* indicate the number of amino acids removed from the original peptide. We have then shown that (10) is true when *ρ* = 0.

If states with identical fluorophore counts are equally probable for all states with *ρ* amino acids removed, it can be shown that all states with equal fluorophore counts are equally probable for all states with *ρ* + 1 amino acids removed. For removal of an amino acid that can’t accept fluorophores under the experimental setup this is trivial, so consider a peptide with *ρ* removed amino acids, an N-terminal amino acid which accepts fluorophores of color 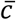, and 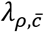 amino acids total which can accept a label of color 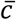. Then let 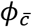 represent the number of remaining functional fluorophores for the peptide, satisfying 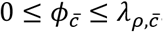.

There are several conditions of the peptide with 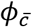 functioning fluorophores scattered among the 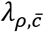 amino acids that can accept a label. When we remove the N-terminal amino acid, we may or may not remove with it a functioning fluorophore. The states which do have a functioning fluorophore in the N-terminal position (only possible when 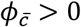) will have their other 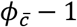 fluorophores distributed between the 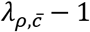 remaining amino acids which can be labeled. Furthermore, these states are equally likely, as they are a subset of the equally likely states with 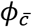 fluorophores. Since trivially 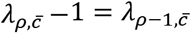, these states map one-to-one with the states for the peptide with one less amino acid remaining, when it has 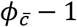 dyes.

When 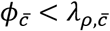, there are states with no fluorophore in the N-terminal position, even though the N-terminal amino acid can accept one. Then the 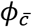 fluorophores will be distributed with equal probabilities among the 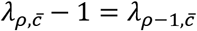 remaining amino acids which can be labeled. Similarly to the other case, these states map one-to-one with the states for the peptide less one amino acid when it has 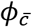 dyes.

The equally distributed probabilities and one-to-one correspondence between states across this amino acid removal ensures that these transformations do not break our guarantees of equal probabilities for *ρ* + 1 amino acids removed. Iteratively applying this reasoning, starting with *ρ* = 0, until we prove that states where *ρ* = *α* are equally likely if they have the same fluorophore counts, demonstrates that (5) is true under the assumptions we have taken.

This proves (10), which allowed us to safely merge states that share both the same fluorophore counts by color and the same numbers of amino acids.

#### Transition probabilities

We also need to know the transition probabilities for our new reduced state space. To deal with peptide detachment is trivial. Dye-loss, either for dyes missing before sequencing begins, or from chemical destruction during sequencing, can be modeled with a binomial distribution. This follows from the assumption that the fluorophores behave independently of each other.

For Edman degradation, there is of course a probability of success or failure of the degradation step, which we model as a Bernoulli random variable. In the case of success, we employ an additional Bernoulli random variable to model the probability of losing or not losing a functioning fluorophore. Because the true states within a merged state are equally likely, we can use combinatorics to count the num er of states which will lose a dye, and the num er that won’t. Together these values can be used to find the probability of losing a fluorophore given a successful Edman degradation, as shown in the following formula, which conveniently reduces to a simple fraction:

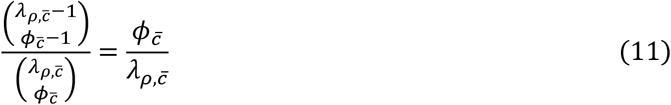

#### State reduction conclusions

This state reduction provides a considerable algorithmic complexity improvement to the HMM forward algorithm. The complexity of the forward algorithm is *O*(*S*^2^*T*), where *S* is the number of states, and *T* is the number of timesteps. Then, if implemented with the true state space of a labeled peptide, the number of states *S* is *O*(*α*2^Λ^), and we get a complexity of *O*(*α*^2^4^Λ^*T*) for the HMM forward algorithm, where *α* is the number of amino acids and Λ is the total number of fluorophores (of any color). However, if we use the reduced state space, then *S* is 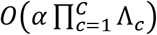, giving an algorithmic complexity of 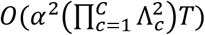 for the forward algorithm, where *C* is the number of fluorophore colors being used and Λ_*c*_ is the number of fluorophores of color *c*. The scaling in either case is dominated by values of Λ or Λ_*c*_, which ranges from 1 to about 25 for human tryptic peptides, though in rare cases Λ_*c*_ can exceed 100.

### A2 – Transition matrix factoring

#### The concept

Multiplication by sparse matrices is far more efficient than with dense matrices. Matrix vector multiplication with a dense matrix is *O*(*S*^2^) where *S* is the size of the vector; for this application vectors with thousands of entries are not uncommon, and even larger vectors are possible, although this depends on the protease and labeling scheme used. For a sparse matrix, matrix vector multiplication can be made to be *O*(*V*), where *V* is the number of non-zero entries in the matrix. For highly sparse matrices this can be a significant improvement.

Since peptides cannot gain amino acids or functioning fluorophores during sequencing, a basic transition matrix for fluorosequencing has zeros except for entries for transitions in which the numbers of fluorophores of each color is decreasing or staying the same. While this does reduce the number of necessary operations, it only does this by a constant factor, with no effect on the asymptotic behavior in the limit. Additionally, the number of amino acids either stays the same, decreases by one (from a successful Edman degradation), or decreases to zero (from a peptide detachment event). This did improve the asymptotic behavior in the number of non-zero entries of the transition matrix, reducing this from 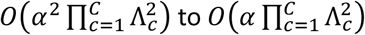.

However, we did better by factoring this matrix (**Figure 4**). We used the independence of our different forms of error, with one matrix in the factored product for each type of error. To demonstrate this factorization, we reformulated our problem in tensor notation. The vector for the state space of a peptide with *C* colors not undergoing Edman degradation or peptide detachment can be viewed as a tensor of order *C*. Each index of the tensor maps to the fluorophore counts of a different color, and the value of an index *i*_*c*_ indicates the number of functioning fluorophores of color *c*, and satisfies 0 ≤ *i*_*c*_ ≤ Λ_*c*_. We also have indices *j*_*c*_ which are similarly defined. Since the transition matrix is a linear mapping from and to this tensor of order *C*, it is necessarily of order 2*C*. We use the Einstein summation convention, and three multi-indices ***i*** = *i*_1_*i*_2_ … *i*_*C*_ and ***j*** = *j*_1_*j*_2_ … *j*_*C*_ and ***k*** = *k*_1_*k*_2_ … *k*_*C*_ for convenience. the matrix vector multiplication operation for one step of the HMM forward algorithm is then given by:

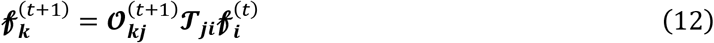

Where (*t*) and (*t* + 1) indicate the timestamp of the values in the order *C* tensor ***ℱ***^(*t*)^, which is indexed by the numbers of working fluorophores for each color and is the tensor form of ***f*** from (1), **ℱ** is the transition matrix ***T*** converted into tensor form, **𝒪** is the emission matrix ***O*** converted into tensor form.

#### Considering fluorophore loss only

Assuming no interactions between different fluorophores and ignoring Edman degradation and peptide detachment, **ℱ** satisfies the following equation:

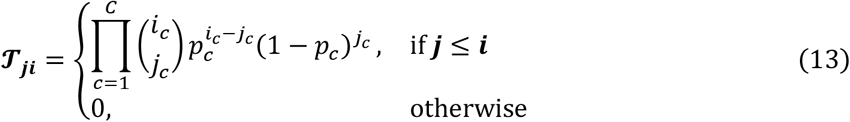

Where *p*_*c*_ is the per cycle dye loss rate of the fluorophores for color *c*. This is simply the product of the binomial distributions for each indexed color of fluorophore. To improve the sparsity of this representation, we can factor **ℱ** into second order tensors **ℬ**^(1)^**ℬ**^(2)^ … **ℬ**^(*C*)^ such that:

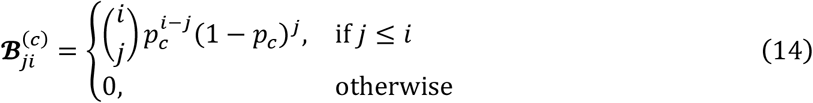

This produces a factorization of **ℱ**:

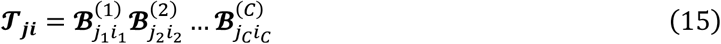

We can plug this into (12) and find:

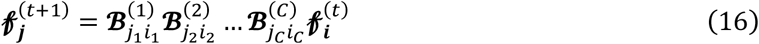

This reduces the algorithmic complexity in this simple case from 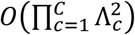 to 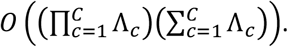

#### Fluorophore loss and Edman degradation

We can expand on this to consider the Edman degradation: In that case we need more indices for the number of remaining amino acids. We modify (12) with additional indices *u* and *v* which satisfy 0 ≤ *u* ≤ *α* and 0 ≤ *v* ≤ *α*, indicating the number of successful amino acid removals, or alternatively the position of an amino acid in the peptide (i. e., the amino acid at the N-terminus of the peptide when *u* amino acids have been removed). This gives:

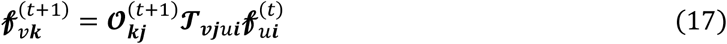

Note that the emission tensor **𝒪** is unaffected by the amino acid count, and depends only on the fluorophore counts, so it does not need to be modified.

We modify **ℱ** from (13) to model Edman degradation, and the exact form of **ℱ** will depend on the peptide under consideration. Let 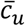 be a number indicating the color of fluorophore at position *u* in the peptide, with a value of 0 indicating no fluorophore, and let 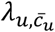 indicate the number of fluorophores of color 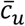 remaining when *u* − 1 amino acids have been removed from the peptide. Then **ℱ** is defined by:

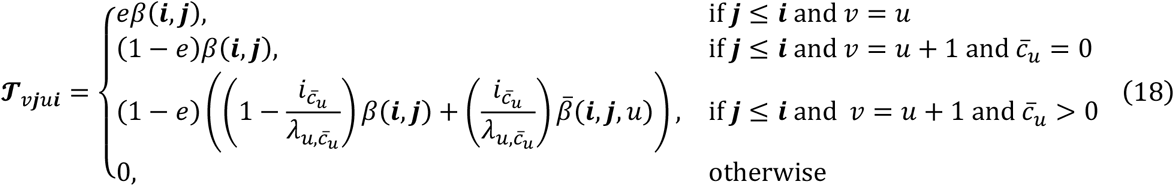

Where:

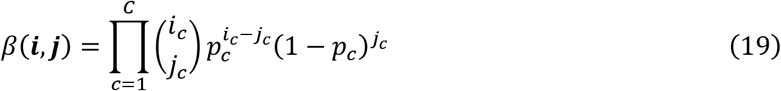

And:

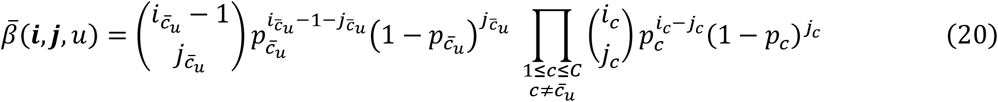

The probability of an Edman degradation failure is essentially the same as in (13), but multiplied by *e* to account for the probability of failure. The probability for a transition involving a successful Edman degradation event which removes an unlabelable amino acid is similarly just like in (13) but multiplied by (1 − *e*), the probability of success. If the amino acid in question is labelable by a color 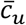, then we may or may not remove a fluorophore of that color in the transition, so we need to take the sum of both possibilities. *β* in (19) gives the standard product of binomials formula from (13), but needs to be multiplied by the probability of no dye loss, which in (18) is 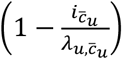. This is then summed with 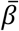 from (20) which gives the product of binomial probabilities starting with one less fluorophore of the color 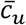, which in (18) is multiplied with the probability of losing a fluorophore with the Edman degradation, 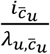. The sum of these two possibilities is then multiplied by the probability of an Edman degradation success, given by (1 − *e*).

To make this more efficient, we introduce a new tensor ***ε*** which represents a transformation for Edman degradation. We define tensor ***ε*** as:

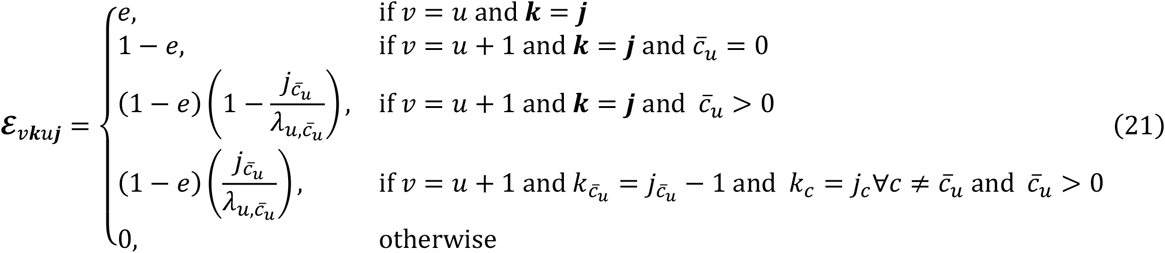

This provides the following factorization of **ℱ**:

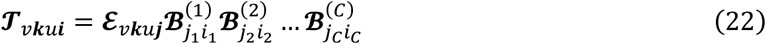

By substituting into (17) and adding an additional multi-index ***l*** = *l*_1_*l*_2_ … *l*_*C*_ we get:

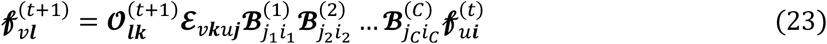

Despite its high dimensionality, ***ε*** is highly sparse, with no more than three non-zero entries per column (here, meaning column in the original non-tensor form matrix). This reduces the algorithmic complexity from 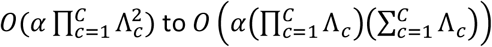. We note that while the extraction of the Edman degradation tensor appears to have little direct effect on the algorithmic complexity reduction, which is because it has a sparsity effect on the original transition tensor, properly handling Edman degradation is critical to this decomposition. We feel this is the easiest way to do this while also factoring the fluorophore loss effects into separate tensors.

#### Everything all together

Handling peptide detachment is simpler. We modify **ℱ** to be:

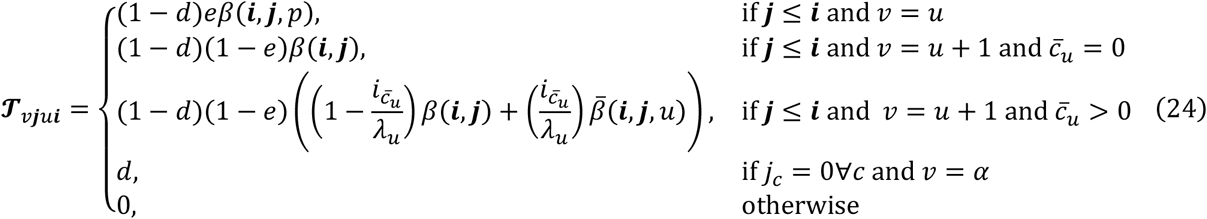

This creates a new “empty” state which can always e transitioned to with pro a ility *d* of detachment. The probability of avoiding this state is (1 − *d*). The functions *β* and 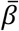 are the same as before in (19) and (20). The matrix vector multiplication step of the HMM forward algorithm has not changed from (17). We can then construct a new tensor **𝒟** for peptide detachment which satisfies:

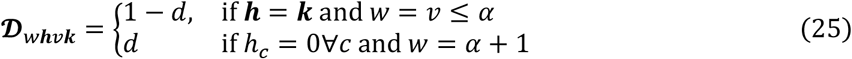

Then we find that:

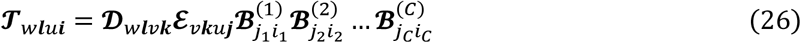

Substituting into (17) with another multi-index ***m*** = *m*_1_*m*_2_ … *m*_*C*_ provides:

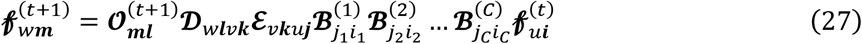

**𝒟** is clearly highly sparse, with two entries in each column of the original matrix in non-tensor form. Thus, **𝒟** has no impact on the algorithmic complexity of this operation. Although **𝒟** and ***ε*** could be combined to achieve this same algorithmic improvement, we found that this separation made our model easier to reason about and work with.

#### Transition matrix factoring conclusions

One of the benefits of this approach to algorithmic complexity reduction is that this factorization provides no loss to the theoretical accuracy of the forward algorithm. No theoretical approximations were necessary, aside from the unavoidable differences in floating-point round-off errors. This allows for highly accurate results with much more efficient runtime characteristics than a naïve implementation.

### A3 – HMM pruning

Because the emission matrix is diagonal, it is equivalent to the diagonal part of its Singular Value Decomposition (SVD), but with a reordering of its indices. This makes sparsification of this matrix equivalent to the Eckart-Young-Mirsky theorem; we can keep the largest *r* values for some chosen value of *r*, and replace the rest of the matrix entries with zeros, having the minimum possible impact on the spectral and Frobenius norms for the chosen value of *r*.

Furthermore, we can propagate this sparsification to the transition matrix. Consider the forward algorithm, with ***T*** representing the transition matrix, and ***O***^(*t*)^ representing the diagonal emission matrix for time *t*. Then if ***f***^(*t*)^ represents the vector of intermediate probabilities at time *t*, we have:

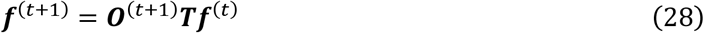

Now we sparsify each ***O***^(*t*)^ as discussed above, to get a series of 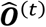. This gives:

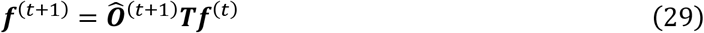

Note that we have many copies of ***T***, which are equal. For our next improvements we need these to be different for each timestep, so we can rewrite (29) with ***T***^(*t*)^ for each timestep *t*, giving

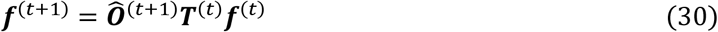

Here the values of many rows and columns of each ***T***^(*t*)^ have been made unnecessary by the sparsification of its neighboring 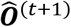 and 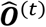, as any vector product with 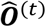will necessarily have zeros except for the *r* entries retained, such that we need only keep the corresponding *r* columns of ***T***^(*t*)^. Similarly, any entry in the vector product with ***T***^(*t*)^ which is not multiplied by one of the *r* entries retained in 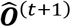 is multiplied by zeros, and is thus unnecessary, so we need only keep the corresponding *r* rows of ***T***^(*t*)^. Calling these approximations 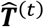, we get

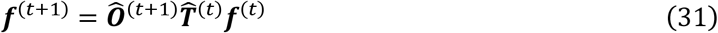

This allows significant sparsity to be used (**Figure 5**). Previously this formula would have been 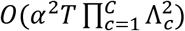 to compute, while this reduces the algorithmic complexity to *O*(*r*^2^*T*). This improvement is beyond what is possible in a more traditional usage of sparse matrix multiplication. For sparse matrix multiplication, we would need to first multiply 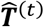 by 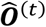 or multiply 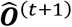 by 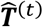. This will only permit you to sparsify your operations on the rows or the columns of ***T*** but not both, giving a complexity of 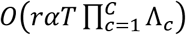. While this is better than not using this inherent sparsity at all, preprocessing the transition matrix in consideration of the emission matrices on either side gives better results in algorithmic complexity. In practice, we use a more complicated pruning scheme, as detailed next.

### A4 – Combining transition matrix factoring with HMM pruning

By making *r* suitably small, HMM pruning can exhibit better algorithmic complexity than if we factor the transition matrix. However, we believe it is much better to combine these algorithmic enhancements (**Figure 6**). To do this, we need to switch into tensor notation, replacing our matrices and vectors with the tensor equivalents we constructed previously. This yields:

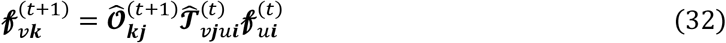

We also want to use the factorization from (26), using timestamp specific sub-tensors of each of the factored pieces. The factorization of (26) becomes:

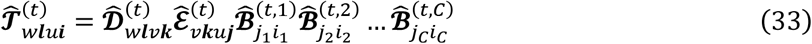

Substituting into (32) gives:

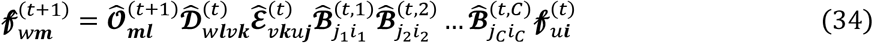

Suppose we were to use standard sparse tensor multiplication techniques and carry this operation out from right to left. Each tensor 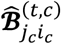 can introduce any entry of input (index *i*) into as many as Λ indices of output (index *j*_*c*_). The resulting computational complexity of the forward algorithm, even with the given sparsity, is then 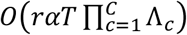.

If we preprocess the computation, pruning each operation now from both directions, the algorithmic complexity does not improve the way it does in the matrix case, although likely this would behave faster in practice. The problem is that the pruning operation itself needs to determine which rows to propagate forwards, which requires accessing every non-zero entry reachable in the forward direction. Many of these values are later pruned in the backwards direction, so the computation itself has much better sparsity, but the time to prune then dominates the algorithmic complexity result.

To improve this further, we add structure to the pruning of 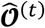. Instead of keeping the *r* largest values in 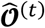, we prune each index of 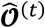 independently. For additional convenience, we limit each index to a contiguous range of values. Then we let each index for any fluorophore color *c* have *r*_*c*_ values and allow 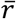 values to index the number of amino acids. These simplifications may cause the pruning to be non-optimal, but we accept this trade-off.

We can then prune our tensors using their known structures (for example, the tensors 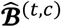 correspond to an upper triangular matrix). This time when we propagate the pruning results in both directions, the time required is only *O*(*C*^2^) (the number of minimum and maximum indices to be propagated through each tensor scales with *C*, as does the number of tensors to be pruned). For the runtime of the tensor operations, consider each tensor individually. 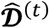 and 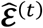are both highly sparse, so they contribute a constant modification to the number of rows or columns when propagating in either direction. 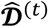 requires special handling. We track the detached state separately from the ordinary range, to avoid unnecessarily includin a lar e ran e of states which don’t need to e.

Then each 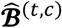 operates on an independent index, and therefore can be considered on its own. This tensor after pruning will have dimensions that are 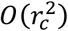, and should have a constant effect on the number of elements input vs output. Therefore, each of these tensors will require an algorithmic complexity of 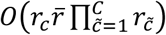. Bringing this all together we get an algorithmic complexity of 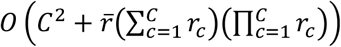 for processing one timestep. The full forward algorithm then has a complexity of 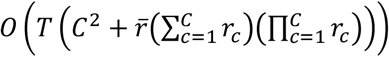.

One remaining clarification is the manner of choosing *r*_*c*_ and 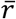. In fact, these values should not be kept constant; let us refer to the values for time *t* as 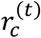 and 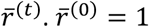 an 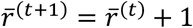, due to the possibility of amino acid removal. These on average are proportional to *α*. To get 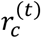, we keep all index values where a fluorophore count of that value has the observed fluorescence intensity for color *c* at time *t* within a specified confidence interval – perhaps within 3*σ* of the mean, where *σ* is the standard deviation of the distribution. These will necessarily be contiguous. The number of indices kept is then 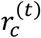.

The standard deviation of a normal distribution scales with the square root of the intensity, and the number of possible index values is limited by the total possible number of fluorophores of color *c*. It follows that any removal of index values proportional to the standard deviation will satisfy 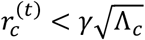 for some constant *γ* dependent on the cutoff. Then the algorithmic complexity is given by 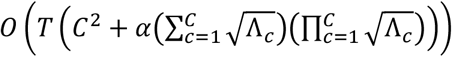.

We chose a specific pruning cut-off by sweeping this parameter and balancing the experimental runtime effects and the precision-recall curves which result from simulated data.

## Appendix B

### Supporting figures

**Figure B1.**
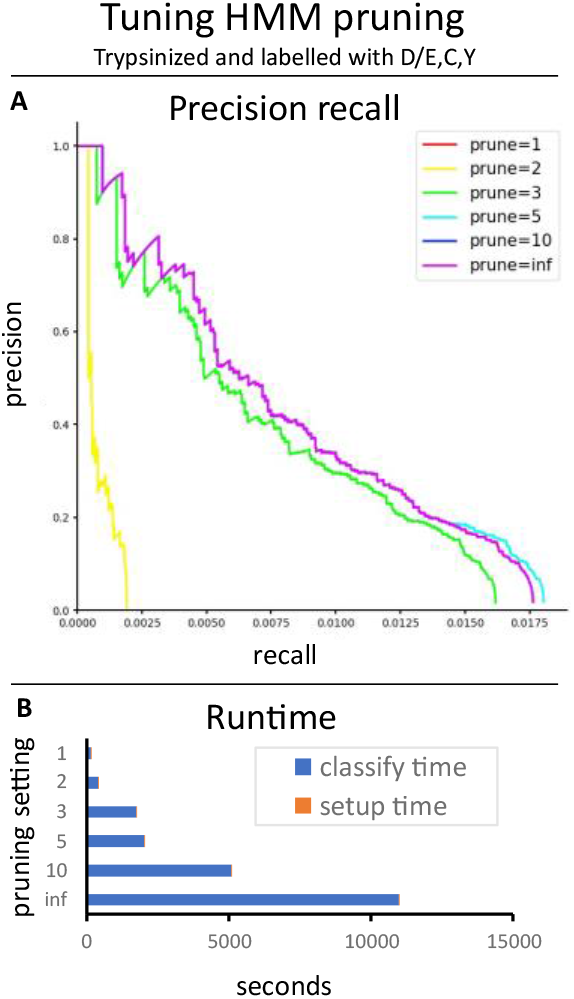
Tuning the pruning parameter of the Bayesian HMM classifier. Setting this parameter to 5 (*i. e*., 5 *σ*) appears to provide the best trade-off. **A:** Precision/recall curves.

**Figure B2.**
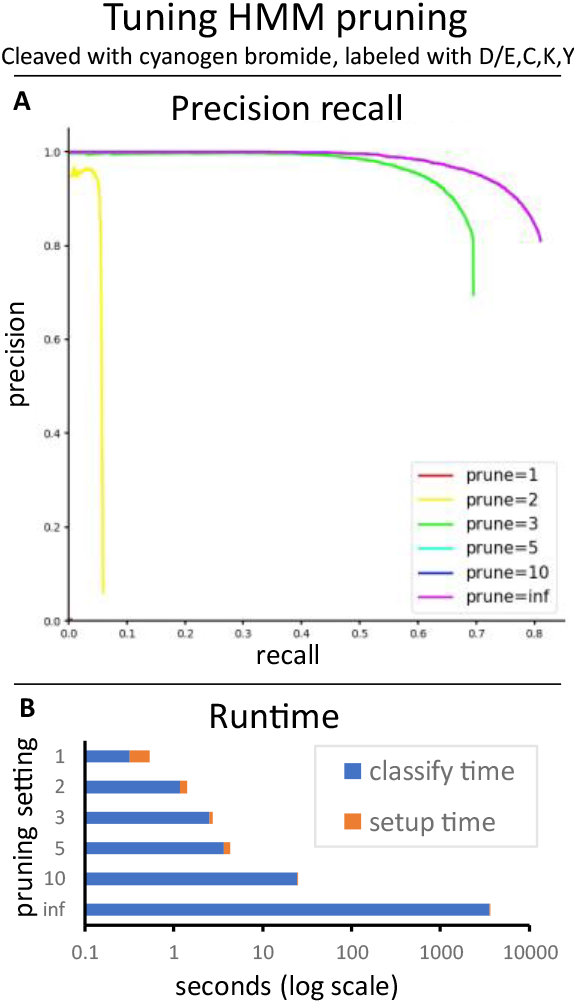
Tuning the pruning parameter of the Bayesian HMM classifier. Here we show a more extraordinary case than **Figure B2. A:** Precision/recall curves. **B:** Runtimes.

**Figure B3.**
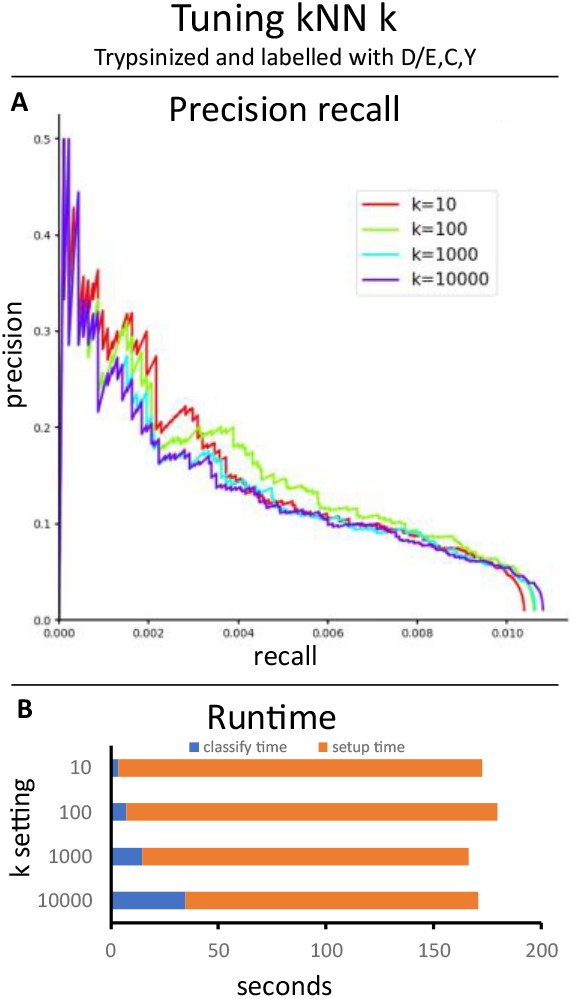
Tuning the k parameter of the kNN classifier. Setting *k* to 10 seems to provide the best trade-off **A:** Precision/recall curves. **B:**

**Figure B4.**
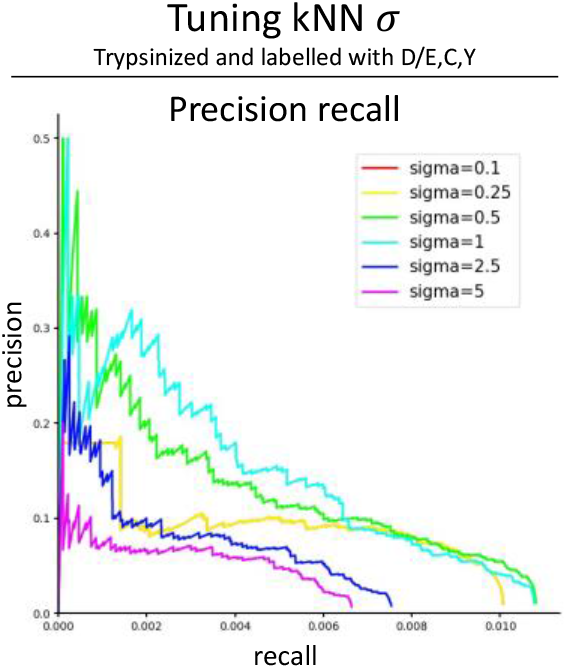
Tuning the *σ* parameter of the kNN classifier. Setting *σ* to 0.5 seems to provide the best trade-off. All settings showed a classify time of about 35 seconds, and a setup time

**Figure B5.**
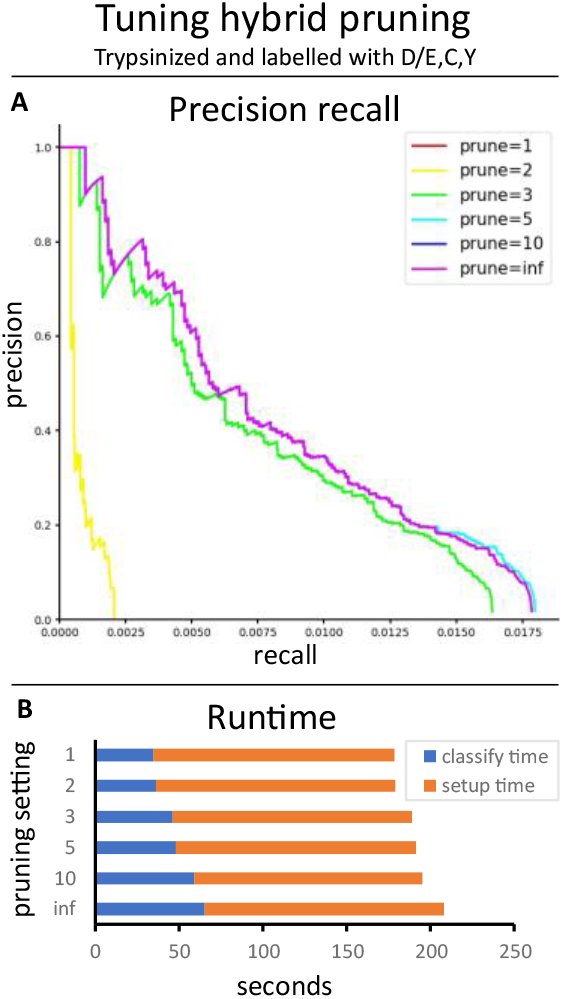
Tuning the pruning parameter of the hybrid classifier. A value of 5 appeared to provide the best trade-off. **A:** Precision/recall curves. **B:** Runtimes.

**Figure B6.**
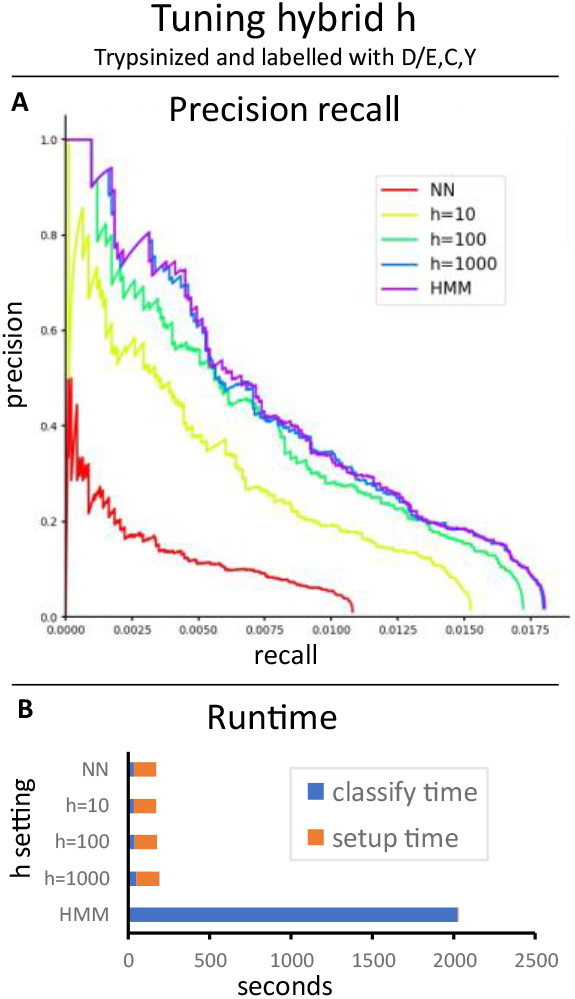
Tuning *h* for the hybrid classifier. An *h* of 1000 provided the best trade-off. **A:** Precision/recall

**Figure B7.**
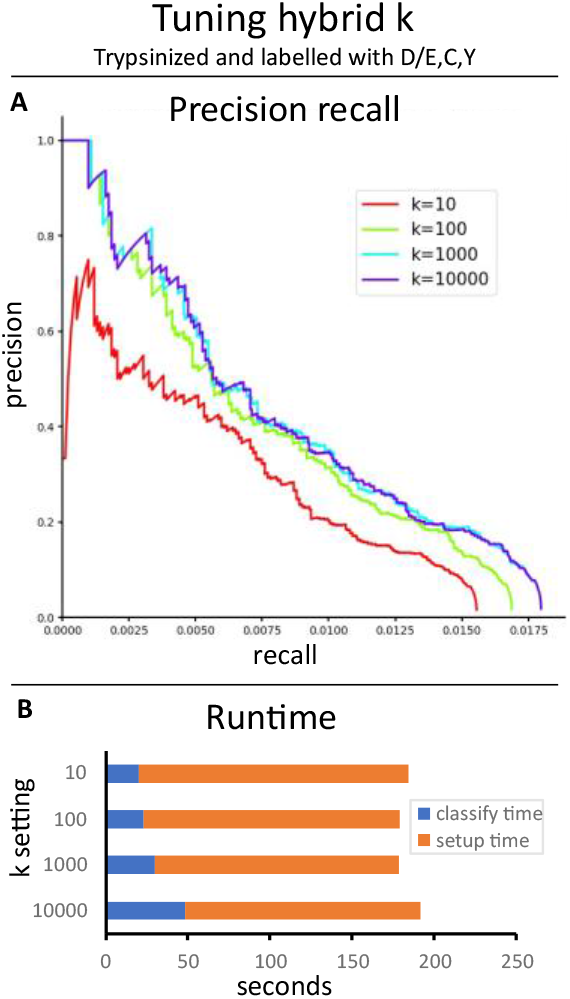
Tuning *k* for the hybrid classifier. A *k* of 1000 or 10000 provided the best results. **A:**

**Figure B8.**
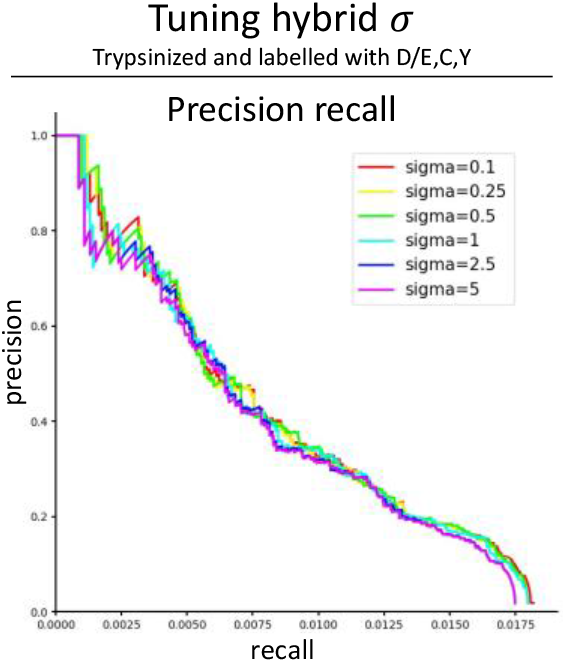
Tuning the *σ*parameter of the hybrid classifier. Setting *σ* to 0.5 seemed to provide the best trade-off. All settings had a classify time of about 50 seconds, and a setup time of about 140 seconds.

